# Quantifying the relative importance of competition, predation, and environmental variation for species coexistence

**DOI:** 10.1101/797704

**Authors:** Lauren G. Shoemaker, Allison K. Barner, Leonora S. Bittleston, Ashley I. Teufel

**Affiliations:** Botany Department, University of Wyoming, Laramie, WY, 82071; Department of Environmental Science, Policy, and Management University of California, Berkeley, Berkeley, CA, 94720; Department of Biology, Colby College, Waterville, ME, 04901; Department of Civil and Environmental Engineering, Massachusetts Institute of Technology, Cambridge, MA, 02139; Department of Biological Sciences, Boise State University, Boise, ID, 83725; Santa Fe Institute, Santa Fe, NM, 87501; Department of Integrative Biology, The University of Texas at Austin, Austin, TX, 78712

**Author notes:** corresponding author, fax: NA. corresponding author:, fax: NA. Data accessibility statement: All code to replicate this study can currently be found on GitHub at https://github.com/lash1937/foodwebs_env_variation_coexistence. Upon acceptance, code will be archived on Zenodo. Statement of authorship: LGS and AKB contributed equally to this manuscript and led the working group from which this project originated. All authors contributed to conceptual development, analyses, manuscript writing, and editing.

**Keywords:** Coexistence theory, ecological networks, species interactions stabilizing mechanisms, environmental fluctuations, diamond model, storage effect

## Abstract

Coexistence theory and food web theory are two cornerstones of the longstanding effort to understand how species coexist. Although competition and predation are known to act simultaneously in communities, theory and empirical study of the two processes continue to be developed independently. Here, we integrate modern coexistence theory and food web theory to simultaneously quantify the relative importance of predation, competition, and environmental fluctuations for species coexistence. We first examine coexistence in a classic multi-trophic model, adding complexity to the food web using a novel machine learning approach. We then apply our framework to a parameterized rocky intertidal food web model, partitioning empirical coexistence dynamics. We find that both environmental fluctuation and variation in predation contribute substantially to species coexistence. Unexpectedly, covariation in these two forces tends to destabilize coexistence, leading to new insights about the role of bottom-up versus top-down forces in both theory and the rocky intertidal ecosystem.

## 2 Introduction

For many decades, community ecologists have developed two complementary theoretical and empirical directions for studying the interactions among species and their dynamic consequences: food web theory (Cohen & Stephens, 1978; McCann, 2011; Terborgh, 2015) and coexistence theory (Chesson, 2000; Barabás *et al.*, 2018). Despite their long independent histories, central to both approaches is a shared interest in explaining the mechanisms that maintain biological diversity, coexistence, and the stability of ecological communities (Ives *et al.*, 2005; Godoy *et al.*, 2018).

Food web theory focuses on consumptive interactions between species across different trophic levels (Fig. 1A), while competition between species is represented through consumer-resource dynamics (e.g., exploitation competition, Godoy *et al.* (2018)). Empirical studies of food webs tend to quantify the presence or absence of links between species at different trophic levels or the frequency of consumptive events among all species in the food web (Berlow *et al.*, 2004; Pascual *et al.*, 2006). From theoretical studies of food webs, we have gained the insight that diverse ecological communities are stabilized by weak interactions between species (McCann *et al.*, 1998), negative feedbacks (May, 1973), and network structures that minimize the frequency of widespread, generalist interactors (McCann, 2011).

**Figure 1:**
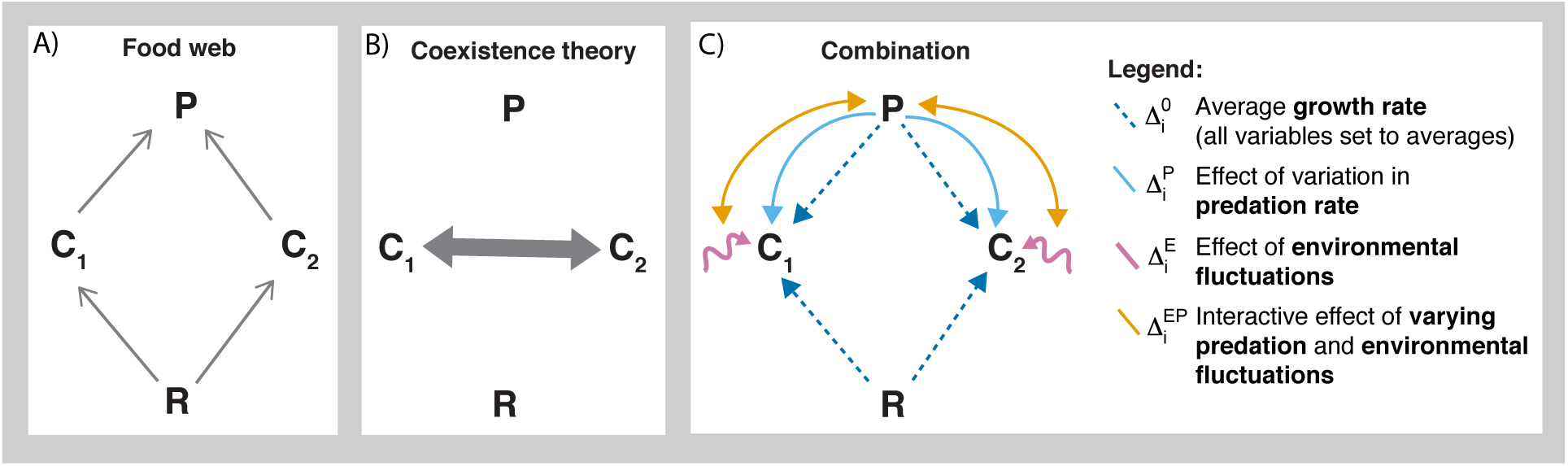
Conceptual diamond model. (A) Food webs generally have links connecting trophic levels in direction of consumption (predator, P, eats consumers, C1 and C2, which share a resource, R). (B) Coexistence theory generally addresses only competition at one trophic level, integrating consumption of a resource into an interaction coefficient. (C) In this framework, we combine food webs and coexistence theory, using the mutual invasion criterion and an approach that allows for coexistence to be partitioned into species’ average growth rates 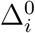, predation variability 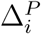, environmental fluctuations 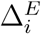, and the interaction between predation and environmental fluctuations 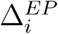.

In comparison, coexistence theory tends to focus on competitive interactions within a single trophic level, exploring how multiple species competing for the same limiting resources, space, or responding to environmental fluctuations can coexist (Fig. 1B). Classic coexistence work shows that diverse and stable communities can occur through three primary mechanisms: (1) the partitioning of resources, where different species are better able to take advantage of different limiting resources, such as nitrogen versus phosphorous in grasslands (Tilman, 1982) (2) trait and demographic trade-offs between species, such as if one species has a higher dispersal rate while another is a superior competitor (Levins & Culver, 1971; Yu & Wilson, 2001), and (3) species partitioning environmental heterogeneity and inherent landscape-level variation, stabilizing overall community dynamics (Chesson, 2000). Current developments in coexistence theory tend to highlight how environmental variation and niche partitioning can promote species coexistence via multiple mechanisms, the most widely studied of which is the storage effect (Chesson, 2000; Snyder & Adler, 2011; Barabás *et al.*, 2018). Empirical applications of this coexistence framework—termed modern coexistence theory (MCT)—tend to quantify demographic rates to examine the relative importance of multiple coexistence mechanisms (Kraft *et al.*, 2015; Germain *et al.*, 2018).

Predation and competition are key forces structuring communities, and at their cores, clear connections exist between these two robust theories for community ecology. Both examine nearly identical questions such as: (1) Why do we observe diverse communities, rather than having one or only a few species dominate? and (2) What properties make diverse communities stable? Past work integrating food webs and coexistence generally fall into three categories: (1) the influence of predators (or “natural enemies”) on the diversity of prey species (Jabot & Bascompte, 2012; Saleem *et al.*, 2012; Griffin *et al.*, 2013), (2) how predator presence alters the strength of competition among prey species (Gurevitch *et al.*, 2000), and (3) coexistence of prey species in the context of predation (Holt, 1984; Holt *et al.*, 1994; Chesson & Kuang, 2008; McPeek, 2019; Klauschies & Gaedke, 2019). Despite the breadth of previous work, few studies have incorporated known food web structure with realistic complexity. The best known example, Brose (2008) implements a consumer-resource model for a diverse simulated food web and analyzes the conditions for persistence in consumers and resources, using an analog of Tilman’s R* theory (Tilman, 1982). While predation has been explicitly incorporated into modern coexistence theory in select scenarios (Chesson & Kuang, 2008; Kuang & Chesson, 2008, 2009, 2010), the approach relies on highly complex underlying analytical derivations, making it difficult to generally apply the framework across more complex theoretical examples, much less empirical systems that require a new system-specific model and corresponding mathematical derivation (Ellner *et al.*, 2019).

Here, we seek to explicitly combine food web theory and coexistence theory, integrating predation and competition together in a conceptual and mathematical framework that is generalizable, easy to use, and can be applied across different systems (Fig. 1C). We extend a recent conceptualization of modern coexistence theory (MCT) (Ellner *et al.*, 2019) to examine how fluctuations in the environment, fluctuations in predator abundance, their interaction, and average fitness differences between consumers can stabilize—or alternatively hinder—coexistence. We first decompose coexistence into its mechanistic components, in a simple but highly studied diamond-shaped food web (McCann *et al.*, 1998; Vasseur & Fox, 2007). Doing so allows us to compare the relative importance of fluctuations in the environment versus predators for the coexistence of two competing species and quantify the strength of top-down versus bottom-up fluctuations. We then examine the generality of coexistence partitioning across parameter space, extending this framework to incorporate additional food web complexity via added consumers and predators. Finally, we apply our approach to a classic empirical ecosystem–the rocky intertidal food web—highlighting its utility in empirical scenarios and insights gained in both theoretical and empirical cases.

## 3 Methods

### 3.1 Partitioning Mechanisms that Promote Coexistence

To examine the relative importance of environmental fluctuations versus predation in maintaining coexistence, we build on the classic framework of MCT (Chesson, 2000). While the framework of MCT traditionally focuses on primary producers—often in annual grasslands (e.g. Mayfield & Stouffer (2017); Kraft *et al.* (2015))—recent advances that allow for a simulation-based approach (Ellner *et al.*, 2019), more easily facilitate application of MCT across a range of applications. Here, we extend the approach of Ellner et al. (2019) to incorporate fluctuations in the environment and predation simultaneously.

In brief, we examine coexistence using the mutual invasibility criterion, incorporating the invader-resident comparison (Barabás *et al.*, 2018). Stable coexistence occurs if each species can successfully invade when rare (i.e. at low-density). Each species’ growth rate when rare, 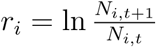, is calculated when the rest of the food web (minus the invader) is at its steady state abundance distribution. To incorporate the invader-resident comparison, we examine average growth rates when rare as 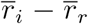 where 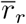 is the average growth rate of resident species within the same trophic level. As the resident species are at steady state abundances and there is a negligible effect of the invader, 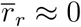, such that 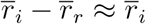. We include the invader-resident comparison, however as it has important implications for the partitioning of coexistence into its mechanistic components, as coexistence can be fostered by factors that either benefit the invader or harm the resident (Ellner *et al.*, 2019; Hallett *et al.*, 2019).

We next decompose each species’ growth rate when rare 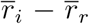 into it’s mechanistic components, following Ellner *et al.* (2019) (See Supplement 1 for the full derivation). The decomposition yields four mechanistic components: the average growth rate with no fluctuations 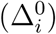, the effect of variation in predator abundance 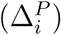, the effects of environmental variation 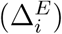, and the interaction effect of both environmental and predator fluctuations 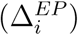 for each species’ growth rate when rare:

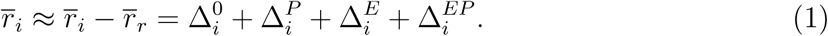

The first term of the decomposition, 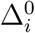, examines species’ ability to invade when rare if both the environment and predator abundances are held constant at their means. We define the second term, 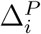, as *nonlinearity in predation*, and it quantifies the effect of variability in predator abundances. If the magnitude of consumptive effects at high predator abundances are less than population gains at low predator abundance (i.e. via saturating consumption), nonlinearity in predator abundance can help stabilize coexistence. Similarly, we define 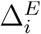 as *nonlinearity in response to the environment*, and it quantifies how underlying variability in environmental conditions can stabilize or destabilize coexistence. Finally, 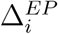 quantifies their interactive effects not accounted for by each main effect in isolation.

### 3.2 Applications of Coexistence Partitioning

#### 3.2.1 Diamond model

Applying the framework above, we first examined the relative importance of environmental fluctuations versus fluctuations in predator abundance using the four-species diamond model (Fig. 1A). This classic model has a long history for analyses of trophic interactions and species coexistence, including in identifying the stabilizing effect of consumer asynchrony in constant environments (McCann *et al.*, 1998), with extensions explicitly incorporating environmental fluctuations (Vasseur & Fox, 2007). Furthermore, in the model, competitors share resources and predators, matching common empirical systems (Williams & Martinez, 2000) in a mathematically simplified and tractable manner.

The diamond model tracks abundance of a predator, *P*, two consumers, *C*_1_ and *C*_2_, and a resource, *R*. Consumer 1 is the superior consumer, but is also the preferred prey species, which maintains coexistence under a variety of parameterizations. In the model, dynamics occur such that:

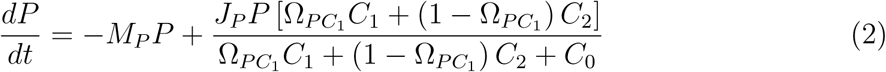

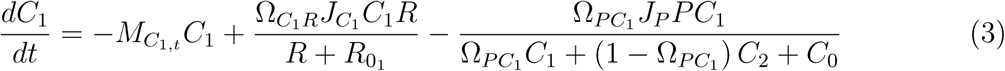

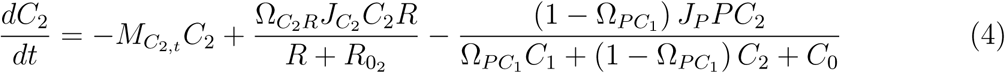

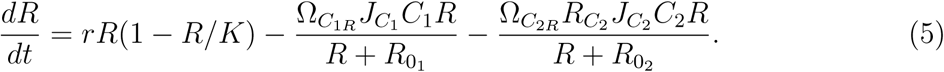

where parameter definitions and values are given in Table 1. Disparities in consumption of the resource and predator preference yields asymptotic dynamics where consumer populations are highly asynchronous and both species co-occur (Vasseur & Fox, 2007).

**Table 1:**
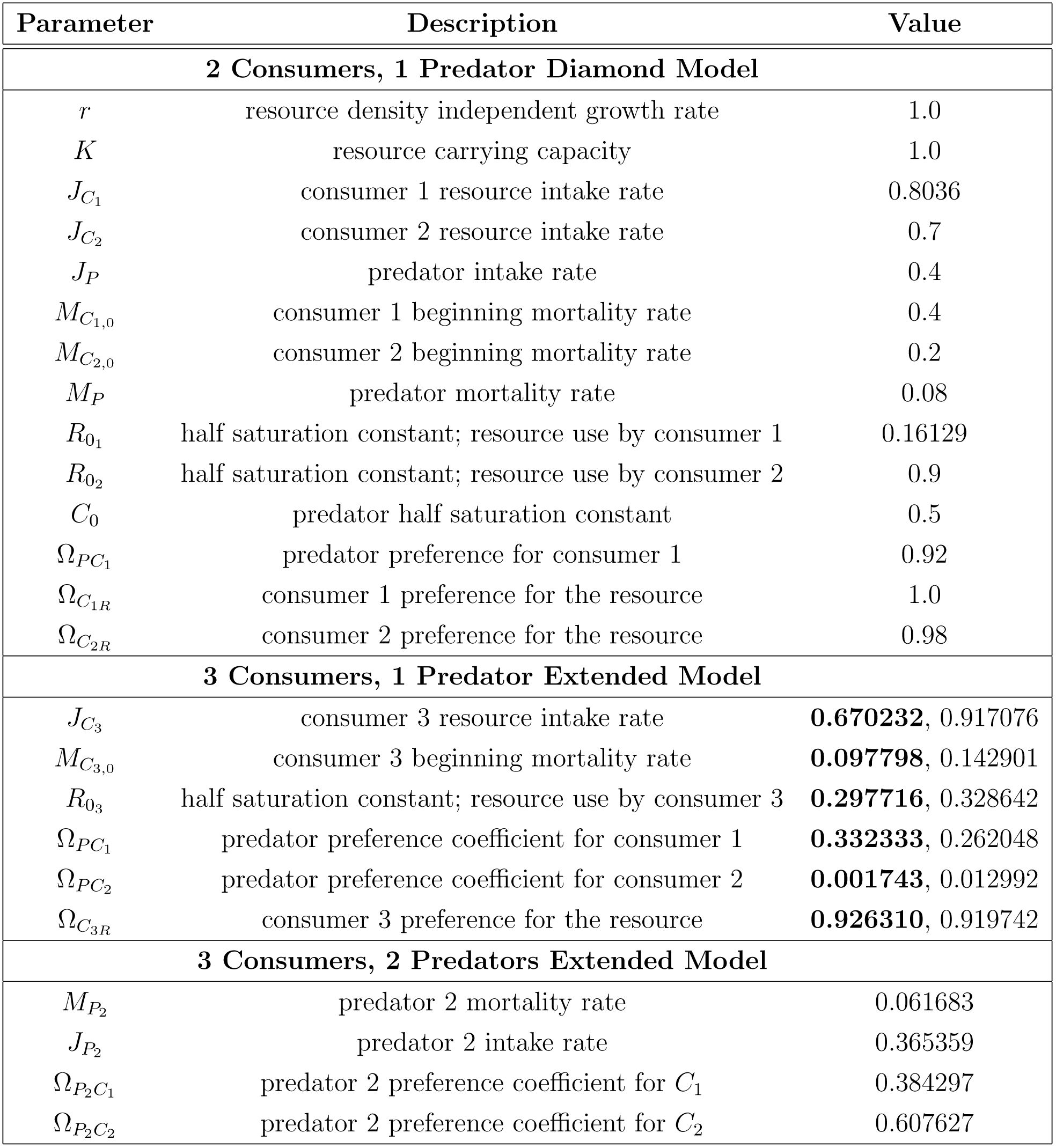
Parameters in the diamond model and its extensions to incorporate additional food web complexity. Parameterization of the classic diamond model matches that from Vasseur & Fox (2007). The mechanistic decomposition of the 2 consumers, 1 predator system is shown in Fig. 3. The bold values from the 3 consumers, 1 predator extended model are those of replicate 1. The mechanistic decomposition of the replicate 1 system are shown in Fig. 5B-D. The values of replicate 2 are given beside the estimates from replicate 1. The mechanistic decomposition of the replicate 2 system are shown in Fig. S3.7D-F. The bold parameters are subsequently used for the 2 predator extension. The mechanistic decomposition of 2 predator extension is shown in Fig. 5F-H.

Environmental variation alters consumer mortality rates, 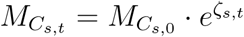, where *ζ*_*s,t*_ are random normally distributed environmental conditions. For each species, *s*, a time series of environmental effects *ζ* is calculated using the Cholesky factorization of the variance-covariance matrix:

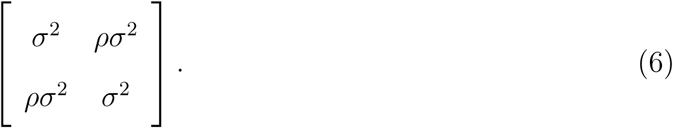

where *σ* is the environmental effect size and *ρ* is the cross-correlation of its effect on consumer species. Multiplying Eq. 6 by a 2 X *T* matrix of random numbers from a normal distribution with mean 0 and unit variance, where *T* is the total number of timesteps to run the model (*T* = 5000), yields *ζ*_1,*t*_ and *ζ*_2,*t*_ for each timestep *t*.

Applying MCT partitioning to the diamond model, we calculated growth rate when rare for each consumer species and its mechanistic decomposition. We set *σ* = 0.55, representative of natural mortality rates from neotropical trees (Condit *et al.*, 1995) and compared scenarios with negative (*ρ* = −0.75), no (*ρ* = 0), and positive (*ρ* = 0.75) cross-correlation. We then examined the relative importance of top-down versus bottom-up controls by varying the predation preference for *C*_1_ from 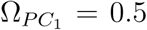 (no preference) to 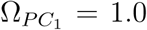 (full preference) in conjunction with varying the strength of environmental fluctuations from *σ* = 0 to *σ* = 0.75. This spans the range of observed environmental fluctuation effects on mortality in both terrestrial and aquatic systems (Vasseur & Fox, 2007; Condit *et al.*, 1995). For each parameter combination, we calculated each species’ growth rate when rare, determining coexistence using the mutual invasibility criterion. Then, using an example with strong predation preference 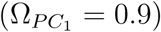, we decomposed coexistence into its mechanistic components for low, intermediate, and high environmental variability (*σ* = 0.1, 0.4 and 0.7 respectively, Supplement 2). All simulations and analyses were conducted in R (R Core Team, 2019).

#### 3.2.2 Expanded diamond model

While the diamond model represents a natural starting-point for examining coexistence, it is a highly simplified food web structure. We therefore expanded the model, first by incorporating a third consumer (*C*_3_) (equation details in Supplement 3, Eq. S3.1). In including a third consumer, we maintained the initial diamond model parameterization except for predator preferences, and applied global optimization by differential evolution to estimate the new parameters 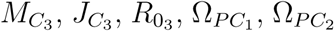, and 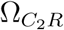 and predator preferences (Table 1) (Ardia *et al.*, 2016). To ensure that populations were stable, we estimated parameters without stochastic mortality of consumers. We created a scoring function based on counting the number of times any of the populations fell below a value of 0.001 over the course of 5000 timesteps (excluding the first 1000 ‘burn-in’ time points) and applied differential evolution to minimize the scoring function using the R package DEoptim, with the DE / best / 1 / bin with jitter (option 3), an initial population size of 100 individuals, and a 0.05 speed of the crossover adaptation (Ardia *et al.*, 2016). We ran the algorithm until the scoring function reached zero and recorded the parameter set from the first member of the population. We repeated this process to compare coexistence mechanisms under two alternative parameter combinations (Table 1). We visually confirmed that these parameter sets resulted in different dynamics in the absence of environmental variation (Fig. S3.1, S3.2) and with environmental variation (Fig. S3.3, S3.4). Using these systems, we calculated coexistence and its mechanistic decomposition when including environmental variation as described above.

To further investigate how complex systems are stabilized, we included a second predator in the model (Eq. S3.2). We followed the same method as above, using the parameter set of replicate 1 from Table 1 to estimate the new parameters for the second predator. We visually confirmed that the estimated parameter set for the model containing two predators resulted in stable dynamics (Fig. S3.5, Fig. S3.6). This methodology allows us to compare coexistence mechanisms with the entire food web (two predators, three consumers, and one resource) and a subset of species from the food web.

#### 3.2.3 Empirical applications in an intertidal food web

Finally, we highlight the applicability of MCT partitioning in empirical systems, focusing on the rocky intertidal ecosystem (Fig. 2), although the framework is applicable across systems. The rocky intertidal communities of the Northeast Pacific Ocean are well-studied model systems for exploring the role of predation and environmental variation in species coexistence (Connell, 1961; Dayton, 1971; Menge *et al.*, 1997; Connolly & Roughgarden, 1999; Forde & Doak, 2004). In this system, a larger barnacle *Balanus glandula* competes for space with the smaller barnacle *Chthamalus dalli/fissus* and with herbivorous limpets (Dayton, 1971). Whelks (predatory snails) and sea stars consume both barnacles, but are not space-limited like barnacles and limpets. Of these five focal taxa, four have planktonic larvae (sea stars, barnacles, limpets), which leads to a decoupling of local population abundance and recruitment (Iwasa & Roughgarden, 1986). Thus, recruitment dynamics are ‘bottomup,’ primarily a function of variation in the environment and the availability of free space (Menge, 2000). Coexistence among sessile, space occupying invertebrates in this system is thought to be controlled both by ‘keystone’ predation (Paine, 1966, 1969) and bottom-up variation in larval supply (Menge *et al.*, 1997), though it remains unclear the degree to which each mechanism contributes to coexistence.

**Figure 2:**
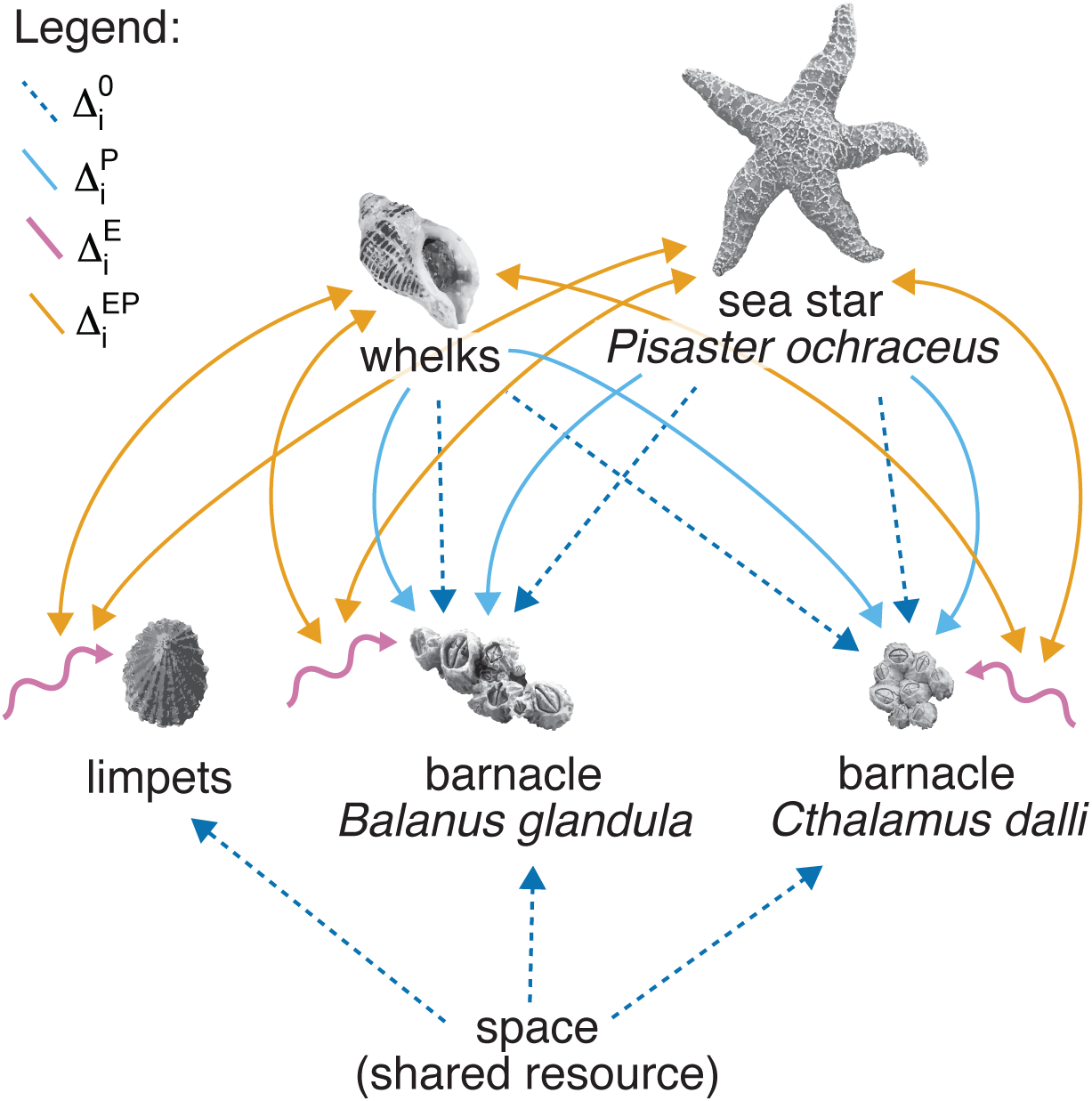
Intertidal food web using conventions introduced in Figure 1C. Barnacles and limpets compete for space, and environmental fluctuations (pink) lead to variation in their larval supply. Sea stars and whelks influence barnacle dynamics through predation (blue). Limpets, however, are not consumed. The lines are not shown for the effect of variation in predation on limpets because the effect is negligible in our model; however, there is a strong indirect, interactive effect of predation and environmental variation (orange).

*Model*. We modeled the rocky intertidal food web (Fig. 2) using a set of stochastic difference equations, based closely on the model and parameterization of Forde & Doak (2004). While here we summarize the model, the full set of equations (Eq. S4.1-S4.10) and parameterization (Table S4.1) can be found in Supplement 4. In brief, the model tracks recruitment dynamics, competition for space, and predator-prey interactions through time. First, the pelagic larval pool for each species (barnacles, limpets, and sea stars) is randomly drawn from a lognormal distribution, based on the range of observed values for the system, mimicking the spatial and temporal recruitment variation in coastal ecosystems (Menge *et al.*, 1997). Next, the realized recruitment from this larval pool to the local ecosystem depends on the availability of free space. Following Shinnen & Navarrete (2014) and Forde & Doak (2004), since no clear competitive hierarchy exists for the rocky intertidal (Menge, 2000), we model competition for free space using lottery competition, where the available free space in the system is calculated at each time step (month) based on the individual size and population abundances of the space-occupiers.

Both recruits and adults of all species are affected by density-independent mortality, while barnacles face additional mortality dependent on predator population size. Predators have the same per-capita effect on both barnacles. In this model, neither prey is preferred over the other (Connolly & Roughgarden, 1999; Forde & Doak, 2004), though predator preference for *Balanus glandula* has been observed (Dayton, 1971) and could be modeled in a future study. Population growth of limpets, sea stars, and whelks is density-dependent, as their predators are not explicitly modeled (Supplement 4).

We simulated the rocky intertidal model 500 times, each for 100 years, tracking population size for each species per month (Forde & Doak, 2004). We examined coexistence across six larval supply scenarios. First, we compared coexistence when all species with pelagic larvae had ‘high’ versus ‘low’ mean supply rates, then subsequently each species was run individually with ‘high’ supply, while all others were held at ‘low’ supply rates (see Table 1 in Forde & Doak, 2004). Coexistence mechanisms were calculated following the procedure described in Section 3.1. Invasion population size was set to 1 for all species, and invasion was simulated after 50 years, such that resident species reached their steady state distribution. Here, variation in the environment manifests as variation in larval supply rather than mortality— as in the diamond model, and nonlinearity in predation, 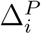, includes variation in predator abundance and recruitment of predators.

## 4 Results

### 4.0.1 Diamond model

When decomposing coexistence into its mechanistic components for the classic diamond model, we find that both consumers are able to stably coexist, as both exhibit positive growth rates when rare, 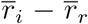 (Fig. 3). For the superior competitor (Fig. 3A), fluctuations in either the environment or the predator abundance matters little for coexistence, as evidenced by the similarity in strength of 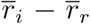 and 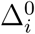, as well as the minimal effects of 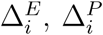, and 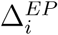. However, for the inferior competitor (Fig. 3B), fluctuations in both predator abundance and the environment increase stability. High predator abundances preferentially increase consumption and decrease the steady state-abundance of the superior competitor, *C*_1_. This decrease in competition between *C*_1_ and *C*_2_ increases the stability of coexistence for the inferior competitor (*C*_2_) via *nonlinearity in predation*. Similarly, *non-linearity in response to the environment* increases stability for the inferior competitor, as positive gains during good years outweigh losses during poor years. However, the interactive effect between environmental fluctuations and fluctuations in predator abundance is destabilizing for *C*_2_. This destabilization of 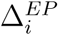 occurs because increased mortality of the resident *C*_1_ decreases consumption and therefore population size of the predator. As *C*_2_ benefits from lower abundances of the superior competitor, the covariance between *C*_1_ mortality and predator abundance destabilizes the growth rate when rare for the inferior competitor, but not enough to hinder coexistence. These results are robust to changes in the cross-correlation of environmental fluctuations (Figs. S2.1, S2.2).

**Figure 3:**
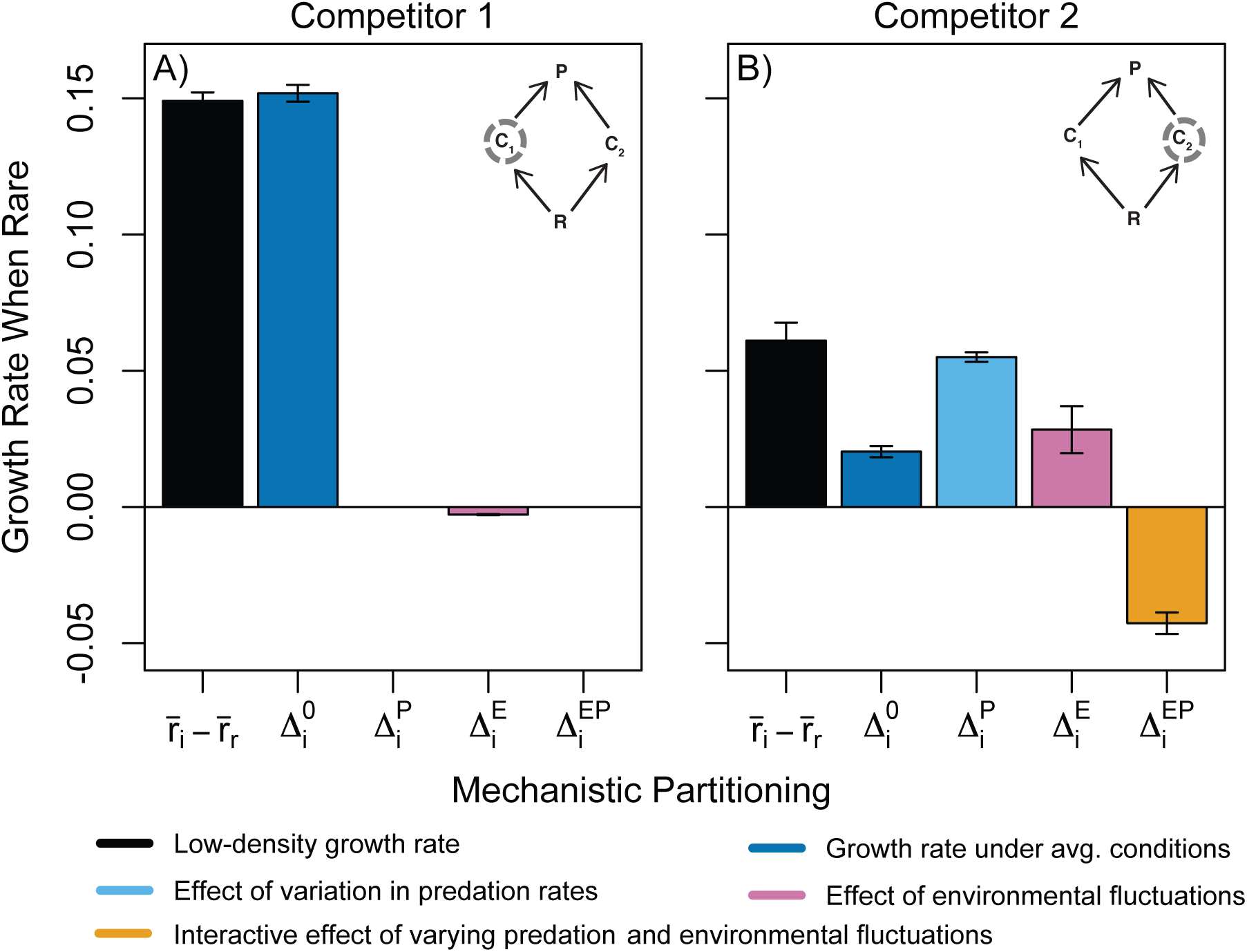
Decomposition of the classic food web diamond model, showing overall growth rate 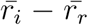, the growth rate with no environmental or predator fluctuations 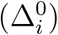, the effect of fluctuations in predator abundance 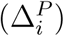, the effect of fluctuations in environmental conditions 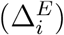, and their combined effects 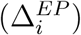. Results are shown for both the superior competitor (A), which is far less effected by variation in either the environment or the predator compared to the inferior competitor (B). Results show mean and standard deviation across 500 runs.

To examine the generality of these results, we calculated both consumer species’ growth rates when rare and coexistence when varying predation preference and the strength of environmental variation. Predation preference had a stronger effect on both coexistence and growth rates when rare compared to the strength of environmental fluctuations (Fig. 4). With no predation preference, *C*_1_ outcompetes the inferior consumer *C*_2_. As predation preference increases, *C*_2_ then is able to outcompete *C*_1_. Only at high preference of the predator for *C*_1_ do both species coexist, as the high predation preference yields oscillatory dynamics that maintain coexistence.

**Figure 4:**
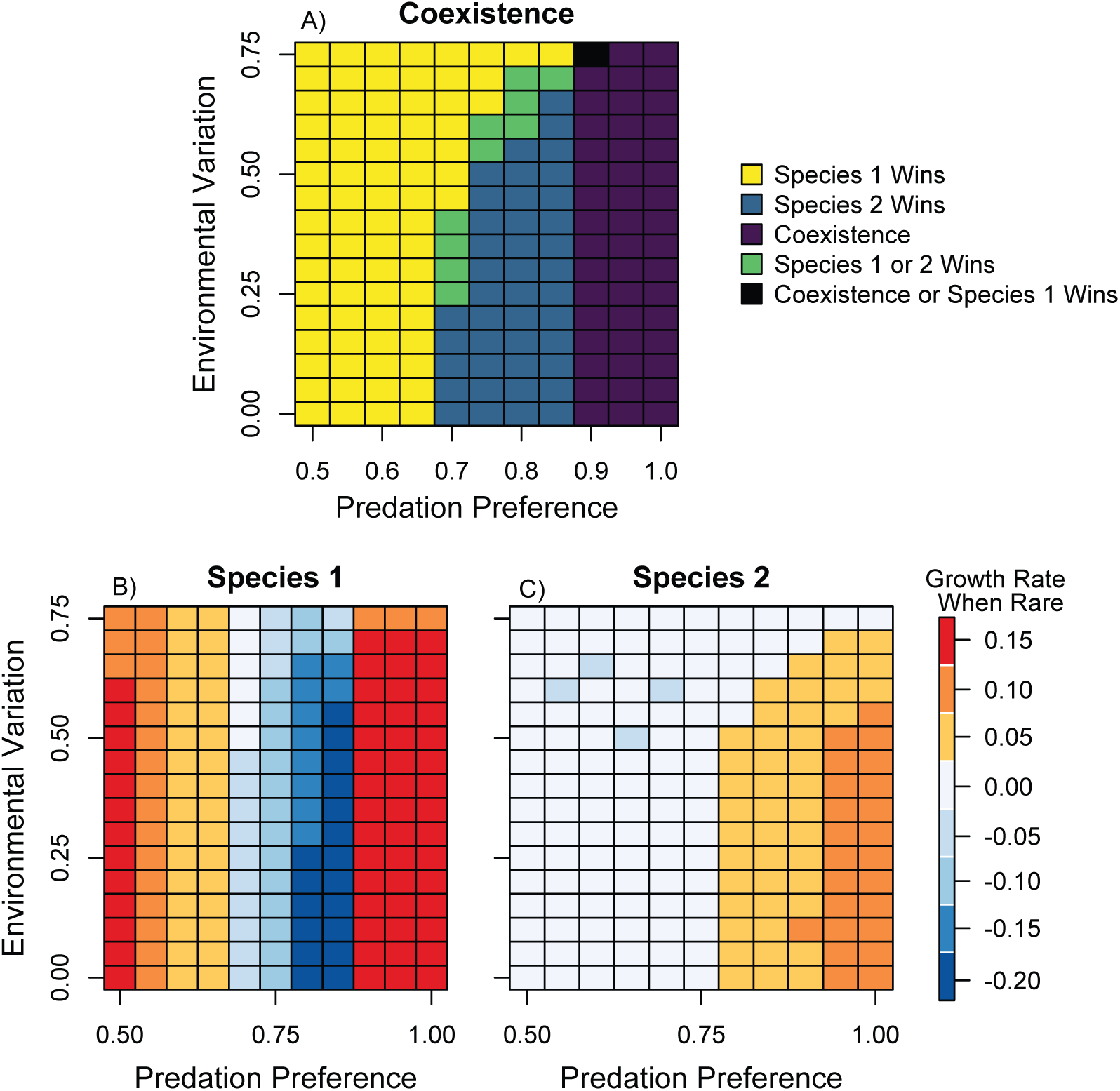
Comparison of the role of predation preference (i.e. top down effects) versus variation in the environment (i.e. bottom up effects) on coexistence of both competitors in the diamond model (A). Results are shown for 1, 760 runs (10 runs per each parameter combination). Coexistence requires that both species’ growth rates when rare are positive (panels B, C; orange and red colors). For species 1, increasing predation preference decreases its growth rate when rare initially, but then allows for coexistence via oscillatory dynamics. For species 2, increasing the predation preference for species 1 increases species 2’s growth rate when rare.

However, while growth rates when rare for each species and overall coexistence depend only moderately on the strength of environmental variation, the relative importance of coexistence mechanisms changes substantially with increased environmental variation (Fig. 1). With low environmental variation, coexistence of *C*_2_ with *C*_1_ is stabilized by 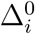 and 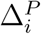. As the strength of environmental variation increases, 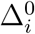 becomes less important and even destabilizing at high environmental variation. Similarly, 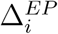 becomes increasingly destabilizing, while 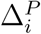 and 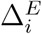 become more stabilizing. The combination of these mechanisms yields coexistence regardless of the amount of environmental variation, *σ*_*ζ*,_ but highlights that the relative importance of coexistence mechanisms changes with increasing environmental variation and that their effects often counteract one another.

### 4.1 Expanded diamond model

To assess how coexistence changes with increasing food web complexity, we first examine adding another competitor to the diamond model (Fig. 5A shows results for the first parameter set). We find that the inclusion of a third competitor causes a reduction in *C*_1_’s growth rate when rare and destabilization by variation in predation and and environmental fluctuations (Fig. 5B). Further, our two parameter sets result in differing stabilizing mechanisms for *C*_2_ (Fig. S3.7), where the effect of 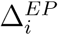 can be stabilizing (Fig. S3.7 B) or destabilizing (Fig. S3.7 E). These findings suggest that the stability of a food web is achieved not just through its structure, but as a function of how the species interact with one another. Furthermore, as the classic diamond model represents a subset of this larger food web (with the same parameterization), comparing Fig. 3 and Fig. 5 highlights the different expectations for coexistence when only considering part of the larger ecological community.

**Figure 5:**
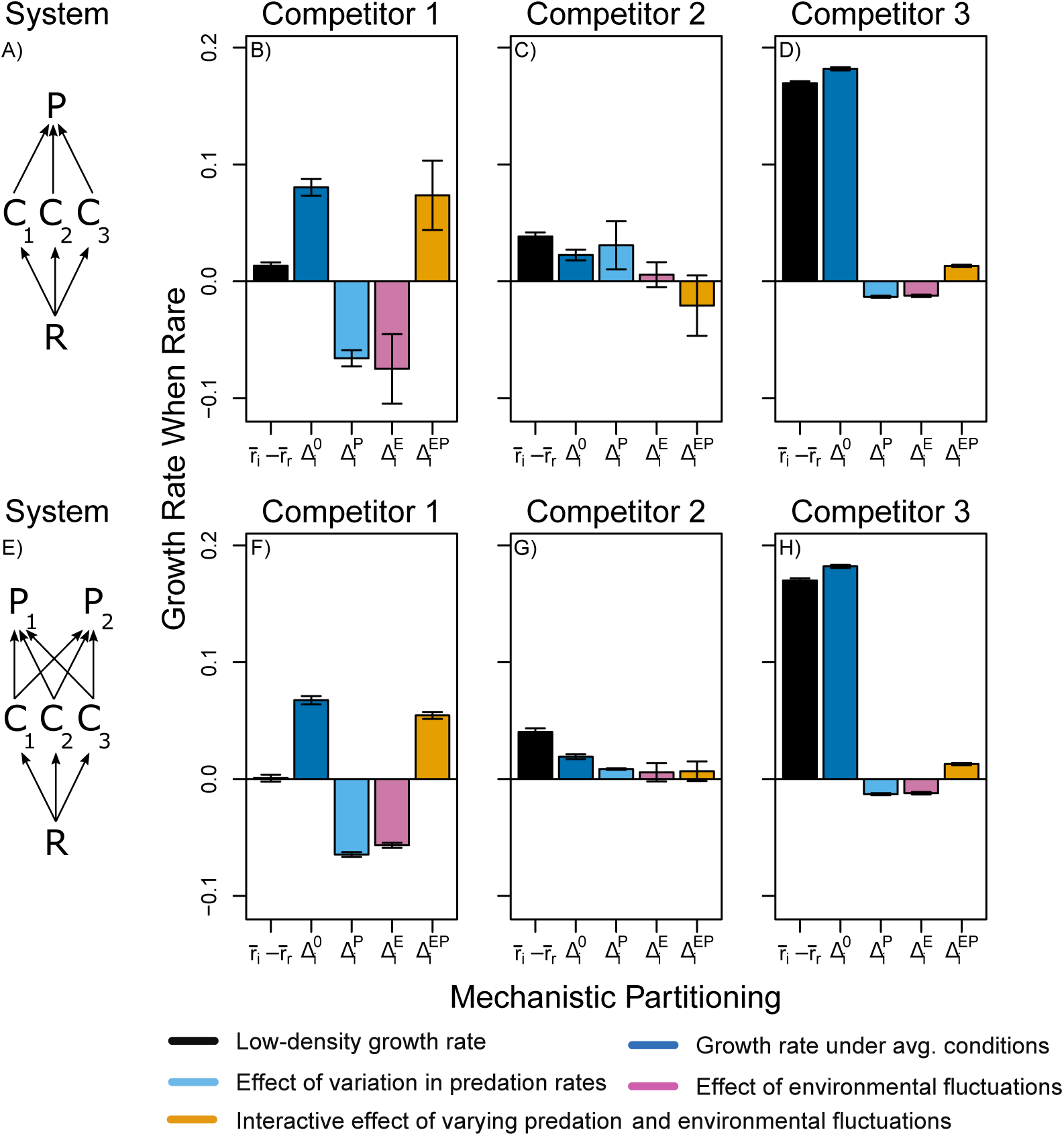
Decomposition of the expanded diamond model, where parameters for Competitors 1 and 2 are the same as in Figure 3 and Competitor 3’s are the bold parameters in Table 1. (A) Illustration of food web with three competitors. (B-D) Mechanistic partitioning for each of the competitors in food web containing three competitors. (E) Illustration of food web with three competitors and two predators. (F-H) Mechanistic partitioning for each of the competitors in food web containing two predators and three competitors. Results of mechanistic partitioning are shown with mean and standard deviation calculated from 500 runs.

To compare how mechanisms of stabilization change when a second predator is included (Fig. 5E) we decompose the coexistence mechanisms of this expanded system (Fig. 5 F-H). The inclusion of a second predator leads to a further reduction in the growth rate when rare of the superior competitor (*C*_1_, Fig. 5F). The second predator also decreases stabilization by *nonlinearity in predation* for *C*_2_ and causes the effect of 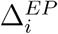 to switch from destabilizing to slightly stabilizing (Fig. 5G). Again, we find different expectations for coexistence strength and its mechanistic comparison when comparing the full model to ones that only consider a subset of species interactions. In aggregate, our results from decomposing increasingly complex food webs demonstrate that the nature of interactions as well the food web topology impact the mechanism by which species coexist.

### 4.2 Empirical applications in an intertidal food web

Finally, we examined coexistence in a temperate rocky intertidal ecosystem, a classically studied system in which both predation and environmental variation have been shown to influence species coexistence. Here, variation in the environment drives variation in larval supply rates of the three taxa that compete for space. In this model, under scenarios where there is no variation in larval supply or predator abundance, the top competitor is the smaller barnacle, *Chthamalus dalli*, though both barnacle species coexist (Fig. 6). When larval supply is low across all planktonic taxa (barnacles, limpets, sea stars), this variation in larval supply benefits both barnacle taxa equally, and *nonlinearity in response to the environment* is the strongest mechanism promoting coexistence. *Nonlinearity in predation* also promotes coexistence of barnacle prey, though to a lesser extent than variation in the environment. Higher positive invasion growth rates under predation suggests a potential ‘hydra effect’ of sea stars and whelks on their barnacle prey (Abrams, 2009). However, coexistence is strongly destabilized by covariation in predator abundance and environmental conditions that promote barnacle growth, paralleling Fig. 3.

**Figure 6:**
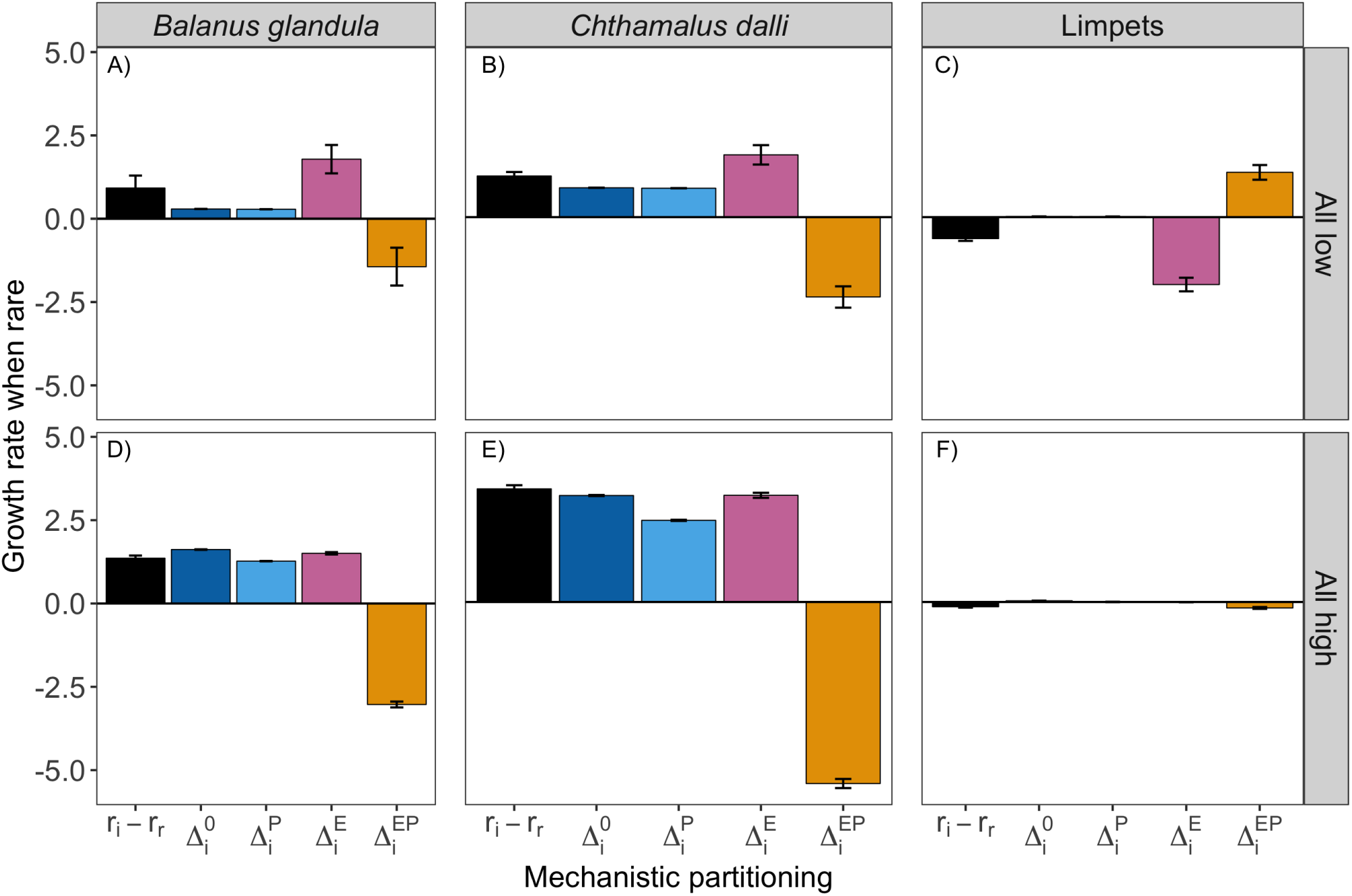
Application of coexistence partitioning to the empirical rocky intertidal foodweb, comparing two levels of larval supply (high and low). Results show the mean and standard error for 500 replicates, each run for 100 years. Both barnacles coexist, despite a destabilizing effect of the interaction between environment and predator fluctuations 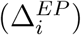. Limpets exhibit a slight negative growth rate when rare, suggesting competitive exclusion. Note that neither predators consume limpets, thus the effect of 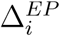 on limpet coexistence strength is an indirect effect mediated by predation on barnacles (see Fig. 2). Additional larval supply scenarios are presented in Fig. S4.1

When larval supply is low, covariation in the environment (larval supply) and predation 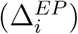 greatly benefits limpets, who have no predators in this model. In other words, limpets benefit from the co-occurrence of high predator abundance and high larval supply, as high predator abundance reduces the abundance of species that compete with limpets, though not enough to allow limpets to ultimately coexist (Fig. 6). In fact, across all scenarios, limpets do not have a positive growth rate when rare (Fig. S4.1). Variation in the environment 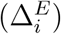, in particular, destabilizes limpet coexistence under low larval supplies enough to counteract the stabilizing effect of 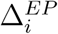.

## 5 Discussion

While coexistence theory and food web theory examine similar core questions, they do so from traditionally disparate perspectives. Connections between the theories are becoming more common (e.g. Kuang & Chesson (2008); Sommers & Chesson (2019)), including recent developments that incorporate niche and fitness differences (Godoy *et al.*, 2018). A unified framework that incorporates direct competition between species together with effects of predation is necessary for gaining a synthetic understanding of how biodiversity is maintained. Our extension of Ellner et al. (2019), builds directly on the framework of modern coexistence theory (Chesson, 2000), including environmental fluctuations through time and space that can maintain coexistence among competitors via niche partitioning (Hallett *et al.*, 2019; Letten *et al.*, 2018). Critically, our extension of MCT also incorporates predation and fluctuations in predator abundances, allowing both bottom-up and top-down mechanisms to be incorporated simultaneously. The relative importance of both—as well as their interactions, as defined by the term 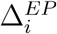—can be examined simultaneously, yielding an extension from the classic framework that allows for the examination of multiple fluctuations across trophic levels. In this study we focus on comparing bottom-up fluctuations in environmental conditions that alter prey mortality rates or larval supply rates and top-down fluctuations in predator abundances and predator larval supply rates.

Applying this approach, it becomes apparent that fluctuations are not always necessary for stabilizing species’ growth rates when rare (e.g. Fig. 3A versus B), but rather individual species may preferentially require fluctuations. These results match insights from modern coexistence theory, even when focusing on a single trophic level in isolation (Hallett *et al.*, 2019; Shoemaker & Melbourne, 2016). Indeed, the superior competitor in the diamond model is only mildly affected by fluctuations in the environment and predator populations, while both of these types of fluctuations increase the stability of the inferior competitor’s (*C*_2_) growth rate when rare. Extending our approach to more complex food web dynamics highlights the importance of considering all key species in a system when quantifying coexistence and corresponding diversity expectations. For example, *C*_1_ exhibited little response to fluctuations in the simple diamond model. Adding a new competitor to the model yielded a destabilizing effect of *nonlinearity in response to the environment* and *predation* (Fig. 5). In general, the interactive effect of environmental fluctuations and predator abundances, 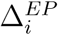, are often destabilizing for the inferior competitor. This may be general across food webs where consumers respond similarly to environmental fluctuations (e.g. Kuang & Chesson (2010)), as environmental conditions that promote consumer growth will correspondingly yield increased predator abundances. Thus, the positive effects of environmental variation may be dampened by increasing predator abundances and therefore realized predation.

More generally, extending modern coexistence theory to examine multi-trophic systems yields key insights into the relative importance of top-down versus bottom-up forces in altering community composition and maintaining biodiversity (Gripenberg & Roslin, 2007; Matson & Hunter, 1992). In many systems, both factors work simultaneously to stabilize community dynamics. Our approach permits the partitioning of fluctuations in bottom-up (e.g. 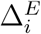) and top-down (e.g 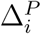) dynamics to examine their relative importance, as well as considering their interaction (e.g. 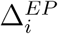). For example, for intertidal barnacle species *Balanus glandula* and *Chthamalus dalli*, both top-down and bottom-up factors stabilize coexistence, although bottom-up factors appear to be slightly more important, at least according to our dynamical model formalization (Forde & Doak, 2004). Empirically, in intertidal systems, the strength of bottom-up factors strongly covary with the strength of top-down processes (Menge *et al.*, 1997). This covariation is destabilizing, as environmental conditions that promote barnacle growth also increase predator abundances. More generally, a similar approach could be applied across ecosystems to partition the relative importance of top-down and bottom-up effects operating simultaneously. For example, in grasslands, nutrient additions often decrease plant biodiversity while herbivores provide a counteracting effect, primarily by reducing light limitation (Borer *et al.*, 2014). Furthermore, in the aquatic detritus-based ecosystems of carnivorous pitcher plants, nutrient additions tend to increase bacterial abundance but not biodiversity, while top predators increase bacterial biodiversity, likely through regulation of intermediate trophic levels (Kneitel & Miller, 2002).

Our approach is general across both theoretical and empirical contexts; however, it requires a tight-coupling of demographic studies with interaction networks to yield dynamical models of both consumer and predator abundances. Measurements of demographic rates and food web interactions are usually made separately—often entire studies in their own rights (e.g. Dibner *et al.* (2019); Gripenberg *et al.* (2019)), and thus all the necessary information is difficult to obtain for many systems. As such, we encourage future work to examine both demographic rates along with trophic links and their corresponding strengths. Doing so will allow for quantifying coexistence and stability, along with the baseline structure of food webs (Pascual *et al.*, 2006). In complex ecosystems, the number of equations and parameters quickly grow with the number of species, so simplifying into functional groups or exploring key species of interest may be necessary for computational tractability.

While here we focus on how the underlying structure of food webs and species’ demography may promote coexistence, a similar approach could additionally incorporate behavioral dynamics. For example, recent work by Sommers and Chesson (2019) show that predator avoidance behavior by prey species can alter coexistence via changes in the importance of apparent competition relative to resource competition. If prey species partition resources, these behavioral changes tend to promote coexistence; when prey species instead partition predators, behavioral modifications undermine coexistence (Sommers & Chesson, 2019). In multiple empirical systems, behavioral changes in prey species via fear-driven avoidance are similarly well documented, such as with brown anoles, green anoles, and curly-tailed lizards on tropical islands (Pringle *et al.*, 2019). Additionally, predators have recently been shown to alter their behavior due to fear-driven avoidance of humans, which positively impacts prey species by increase their foraging ability (Suraci *et al.*, 2019). Partitioning the mechanistic effects of such behavioral changes—including with multiple predator trophic levels—would yield insight into the potential stabilizing effects of behavior on coexistence of prey species.

A further extension of this work could be examining coexistence under global change, with directional changes in environmental fluctuation (Usinowicz & Levine, 2018). For instance, as temperature increases so do metabolic and encounter rates, which likely have important ramifications for coexistence, and in particular the contribution of *relative non-linearity in predation*, 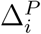 (Moya-Laraño *et al.*, 2014). Additionally, this framework could be extended to examine eco-evolutionary dynamics. Classic single-trophic level applications of modern coexistence theory show the ability for stabilizing coexistence mechanisms to evolve, especially if one species has a greater evolutionary ability (Snyder & Adler, 2011). In contrast, multiple species can co-evolve in a manner that erodes the importance of stabilizing coexistence mechanisms (Snyder & Adler, 2011). Extensions examining phenotypic variation (Gibert & DeLong, 2017) or eco-evolutionary dynamics in food webs may be critical, as both can drastically alter food web dynamics, links, and species’ abundances, even under relatively small variation or selective pressures (Gibert & Yeakel, 2019). Applications to eco-evolutionary food webs could provide insight into the evolutionary and environmental factors impacting species coexistence and community stability.

## 6 Acknowledgements

We thank Artemy Kolchinsky, Andy Rominger, and Margaret Siple for conversations in a Santa Fe Institute (SFI) Research Jam that led to the development of this project. This work was supported by a James S. McDonnell Foundation (JSMF)-SFI Postdoctoral Working Group grant to LGS and AKB (grant number: 357(14)). LGS, AKB, and LSB were supported by JSMF Postdoctoral Fellowships, grants #220020513, #220020478, and #220020477. AIT was supported by the Omidyar Program at the SFI.

## 7 Supporting Information

Appendix S1: Derivation of Coexistence Mechanisms

Appendix S2: The Classic Diamond Model

Appendix S3: Expanding the Diamond Model for Additional Complexity

Appendix S4: Rocky Intertidal Food Web Case Study

