## Supplement 1 for "Quantifying the relative importance of competition, predation, and environmental variation for species coexistence"

#### Derivation of Coexistence Mechanisms: Supplement 1

Lauren G. Shoemaker, Allison K. Barner, Leonora S. Bittleston, and Ashley I. Teufel

#### 1 Derivation

Following classic MCT, we test for stable coexistence using the mutual invasion criterion, which requires that each species can invade when all other species in the food web (minus the invader) are at their steady state abundance distributions. We apply the invader-resident comparison, where we examine the invader's growth rate when rare, subtracting out the resident-consumers' average growth rates (Barabás *et al.*, 2018; Ellner *et al.*, 2019). In this approach, $\overline{r}_i - \overline{r}_r \approx \overline{r}_i$ (where $i$ denotes the invader and $r$ denotes the resident) since the residents' average growth rates will be 0 as they are at their steady state distributions and the invader is at such low density that interspecific competition is minimal. We incorporate the invader-resident comparison, however, as coexistence can be fostered by mechanisms that either help the invader or hinder the resident. Stable coexistence requires that $\overline{r}_i - \overline{r}_r > 0$ for all species, where $r_j = \ln \frac{N_{j,t+1}}{N_{j,t}}$ for a species $j$.

The average population growth rate of each species, $j$, through time depends on both environmental fluctuations ($E(t)$) and fluctuations in predator abundances ($P(t)$), such that

$$\overline{r}_j = \frac{1}{T}\sum_{t=1}^{T} r_j(E(t), P(t)). \tag{S1.1}$$

A population increases if $\overline{r}_j > 0$. Critically, this formalization allows us to decompose growth rates into their mechanistic components. Following Ellner et al. 2019, we for each species we define the following terms:

$$\epsilon_j^0 = r_j(\overline{E}, \overline{P}), \tag{S1.2}$$

$$\epsilon_j^E = r_j(E(t), \overline{P}) - \epsilon_j^0, \tag{S1.3}$$

$$\epsilon_j^P = r_j(\overline{E}, P(t)) - \epsilon_j^0 \tag{S1.4}$$

and

$$\epsilon_j^{EP} = r_j(E(t), P(t)) - \left[\epsilon_j^0 + \epsilon_j^E + \epsilon_j^P\right]. \tag{S1.5}$$

$\epsilon_j^0$ is the population growth rate when the environment and predator abundances are constant at their means, $\epsilon_j^E$ is the main effect of the environment varying around its mean, $\epsilon_j^P$ is the main effect of predator abundance varying around its mean, and the term $\epsilon_j^{EP}$ accounts for the fact that having variability in both the environment and predator abundances will not equal the sum of the main effects.

Following Ellner et al. (2019), we find that

$$r_j(E(t), P(t)) = \epsilon_j^0 + \epsilon_j^E(E) + \epsilon_j^P(P) + \epsilon_j^{EP}(E, P) \tag{S1.6}$$

which we can average to determine that:

$$\overline{r}_j = \epsilon_j^0 + \overline{\epsilon}_j^E + \overline{\epsilon}_j^P + \overline{\epsilon}_j^{EP}. \tag{S1.7}$$

we use Equation S1.7 to compute the invader-resident comparison, where we compare $\overline{r}_i - \overline{r}$. For the invader-resident comparison,

$$\overline{r}_i = \overline{r}_i - \overline{r}_r = \Delta_i^0 + \Delta_i^E + \Delta_i^P + \Delta_i^{EP}. \quad \text{(S1.8)}$$

where $\Delta_i$ is the invader-resident difference between corresponding terms. For example, $\Delta_i^0 = \overline{\epsilon}_i^0 - \overline{\epsilon}_r^0$. The full derivation with further explanation can be found in Ellner et al. 2019. For scenarios with multiple resident consumer species (i.e. the expanded diamond model and the intertidal food web model), we weight all residents equally. For example, $\Delta_i^0 = \overline{\epsilon}_i^0 - \dfrac{1}{S-1}\sum_{r\neq i}\overline{\epsilon}_r^0$, following the general approach of Ellner et al. (2019).

Equation S1.8 provides the full decomposition used throughout this study. Here, $\Delta_i^0$, examines species' ability to invade when rare if both the environment and predator abundances are constant at their means. The second term, $\Delta_i^E$, is *nonlinearity in response to the environment*, and it quantifies the effect of variability in environmental conditions. This directly affects consumer mortality rates in the classic diamond model and the expanded diamond model and larval supply rates in the rocky intertidal food web model. Intuitively, *nonlinearity in response to the environment* can stabilize a species' growth rate when rare if the positive effects of "good" environmental years are larger in magnitude than the negative effects of "bad" environmental years. Similarly, $\Delta_i^P$ is *nonlinearity in predation* and is stabilizing if the magnitude of consumptive effects at high predator abundances are less than population gains at low predator abundance (i.e. via saturating consumption). Finally, $\Delta_i^{EP}$ quantifies their interactive effects not accounted for by each main effect in isolation.

### 2 Applications

We first apply the above decomposition to the classic diamond model and the expanded diamond model. When calculating each mechanism, we ran the resident community for $5000$ time steps. We used the first $2500$ as a "burn-in" to remove any potential effect of starting conditions and defined the last $2500$ time steps as the steady-state distribution. For each time point $t$ in the last $2500$ time steps, we calculated the invader's growth rate when rare and the resident consumers' growth rates with an invader abundance of $0.001$.

For the rocky intertidal food web, we ran the resident community for $100$ years ($1200$ time steps in the model). Similarly to the diamond model, we used the first half as a "burn-in" to remove potential effects of starting conditions and defined the last $50$ years as the steady-state distribution. We calculated the mutual invasion criterion with an invader abundance of $1$ individual.

All code is available in our supplementary material.
