## Supplement 2 for "Quantifying the relative importance of competition, predation, and environmental variation for species coexistence"

### The Classic Diamond Model: Supplement 2

Lauren G. Shoemaker, Allison K. Barner, Leonora S. Bittleston, and Ashley I. Teufel

### 1 Altering Cross-Correlation of Environmental Variation and its Strength

While altering the cross-correlation of environmental variation between species ($\rho_\zeta$) had little effect on the fluctuation dependent coexistence mechanisms (Figs. S2.1, S2.2), increasing the strength of environmental fluctuations ($\sigma_\zeta$) significantly changed the contributions of multiple coexistence mechanisms (Fig. 1). Increasing the strength of environmental fluctuations caused the strength of the growth rate when rare to decrease for the inferior competitor ($C_2$). However, both *nonlinearity in environmental responses* and *nonlinearity in predation* increased, further stabilizing coexistence with increasing environmental fluctuations. In comparison, their interactive effect became more destabilizing (i.e. negative). The superior competitor, however, was only minimally affected by changes in environmental fluctuations ($\sigma_\zeta$).

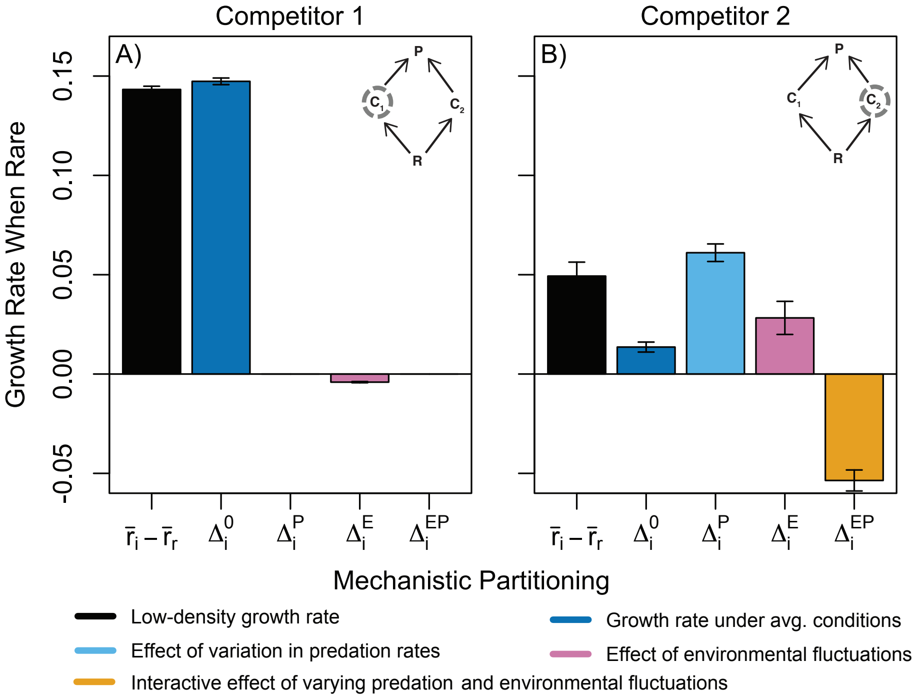


Figure S2.1: Positive cross correlation comparison to Fig. 3 in the main text. Examining the role of predation preference (i.e. top down effects) versus variation in the environment (i.e. bottom up effects) on coexistence of both competitors in the diamond model. Coexistence requires that both species' low-density growth rates are positive (panels A, B; black). Model parameters are the same as in Table 1, except with positive cross-correlation of environmental fluctuations between species ($\rho_\zeta = 0.75$). Results are shown for 500 runs with error bars denoting standard deviation.

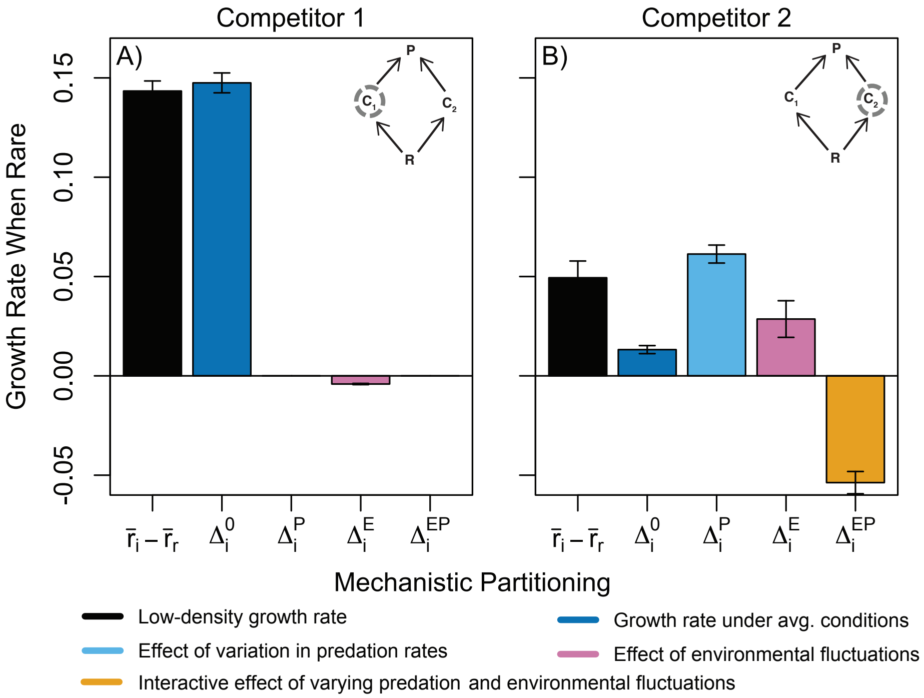


Figure S2.2: Negative cross correlation comparison to Fig. 3 in the main text. Examining the role of predation preference (i.e. top down effects) versus variation in the environment (i.e. bottom up effects) on coexistence of both competitors in the diamond model. Coexistence requires that both species' low-density growth rates are positive (panels A, B; black). Model parameters are the same as in Table 1, except with positive cross-correlation of environmental fluctuations between species ($\rho_\zeta = -0.75$). Results are shown for 500 runs with error bars denoting standard deviation.

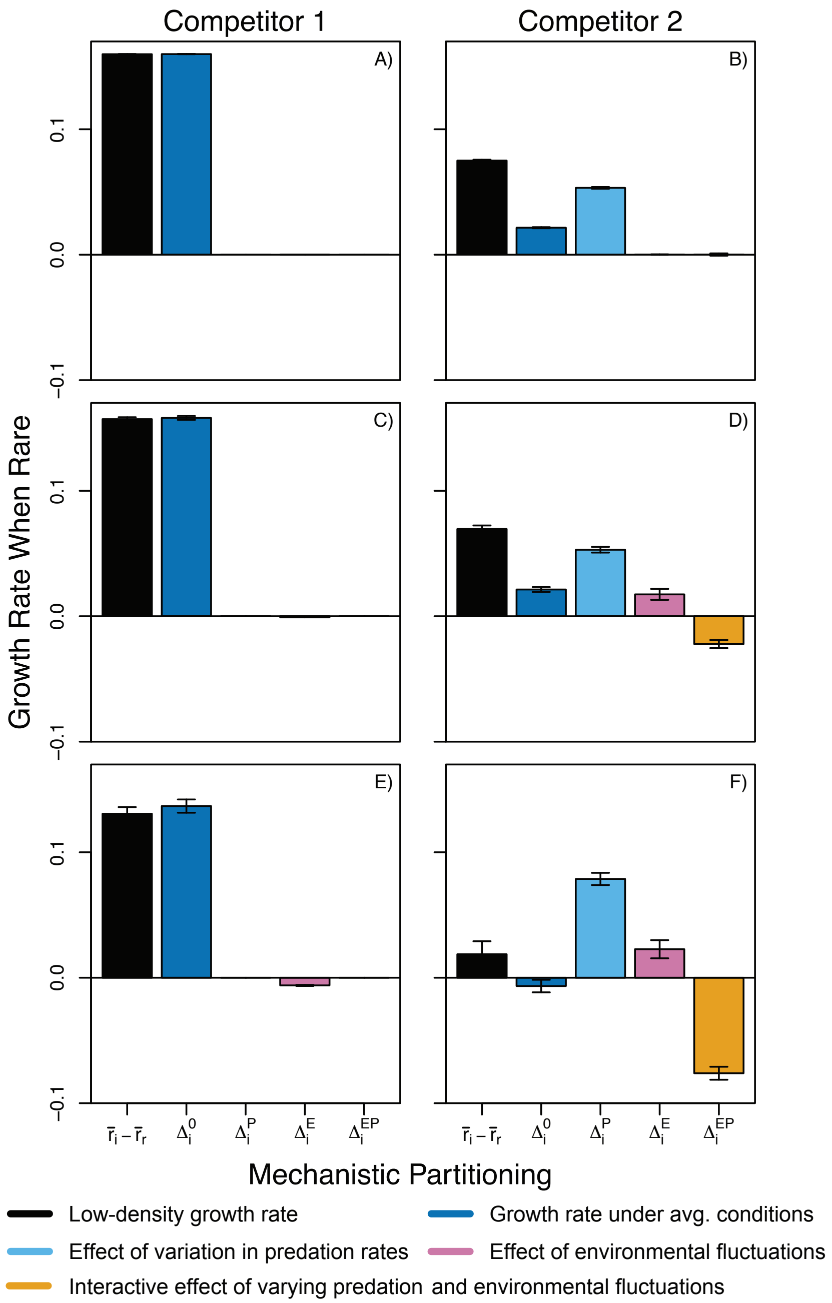


Figure S2.3: Increasing the strength of environmental variation ($\sigma_\zeta$) on coexistence mechanisms. The top row (A, B) shows low environmental variation ($\sigma_\zeta = 0.1$), while the middle row (C, D) shows medium environmental variation ($\sigma_\zeta = 0.4$), and the last row (E, F) shows high environmental variation ($\sigma_\zeta = 0.7$). Each subpanel shows results from 500 runs, where error bars denote standard deviation.
