## Supplement 3 for "Quantifying the relative importance of competition, predation, and environmental variation for species coexistence"

#### Expanding the Diamond Model for Additional Complexity: Supplement 3

Lauren G. Shoemaker, Allison K. Barner, Leonora S. Bittleston, and Ashley I. Teufel

#### 1 Three Consumers, One Predator

Expanding on the diamond model to include a third competitor yields the following set of equations:

$$\frac{dP}{dt} = -M_p P + \frac{J_p P[\Omega_{PC_1} C_1 + \Omega_{PC_2} C_2 + (1 - (\Omega_{PC_1} + \Omega_{PC_2}))C_3]}{\Omega_{PC_1} C_1 + \Omega_{PC_2} C_2 + (1 - (\Omega_{PC_1} + \Omega_{PC_2}))C_3 + C_0}$$

$$\frac{dC_1}{dt} = -M_{C_1} C_1 + \frac{\Omega_{C_1 R} J_{C_1} C_1 R}{R + R_{0_1}} - \frac{\Omega_{PC_1} J_p P C_1}{\Omega_{PC_1} C_1 + \Omega_{PC_2} C_2 + (1 - (\Omega_{PC_1} + \Omega_{PC_2}))C_3 + C_0}$$

$$\frac{dC_2}{dt} = -M_{C_2} C_2 + \frac{\Omega_{C_2 R} J_{C_2} C_2 R}{R + R_{0_2}} - \frac{\Omega_{PC_2} J_p P C_2}{\Omega_{PC_1} C_1 + \Omega_{PC_2} C_2 + (1 - (\Omega_{PC_1} + \Omega_{PC_2}))C_3 + C_0}$$

$$\frac{dC_3}{dt} = -M_{C_3} C_3 + \frac{\Omega_{C_3 R} J_{C_3} C_3 R}{R + R_{0_3}} - \frac{(1 - (\Omega_{PC_1} + \Omega_{PC_2}))J_p P C_3}{\Omega_{PC_1} C_1 + \Omega_{PC_2} C_2 + (1 - (\Omega_{PC_1} + \Omega_{PC_2}))C_3 + C_0}$$

$$\frac{dR}{dt} = rR(1 - R/K) - \frac{\Omega_{C_1 R} J_{C_1} C_1 R}{R + R_{0_1}} - \frac{\Omega_{C_2 R} J_{C_2} C_2 R}{R + R_{0_2}} - \frac{\Omega_{C_3 R} J_{C_3} C_3 R}{R + R_{0_3}}$$

$$\textit{where } (\Omega_{PC_1} + \Omega_{PC_2}) <= 1 \qquad \text{(S3.1)}$$

where parameter values are given in the main text (Table 1). We estimated parameters of this system twice, allowing us to compare coexistence under the same food web structure, but with different interaction strengths.

To confirm that the estimated parameters result in stable dynamics we examine the dynamics of both of these systems without variance (Fig. S3.1, S3.2) and with variance (Fig. S3.3, S3.4).

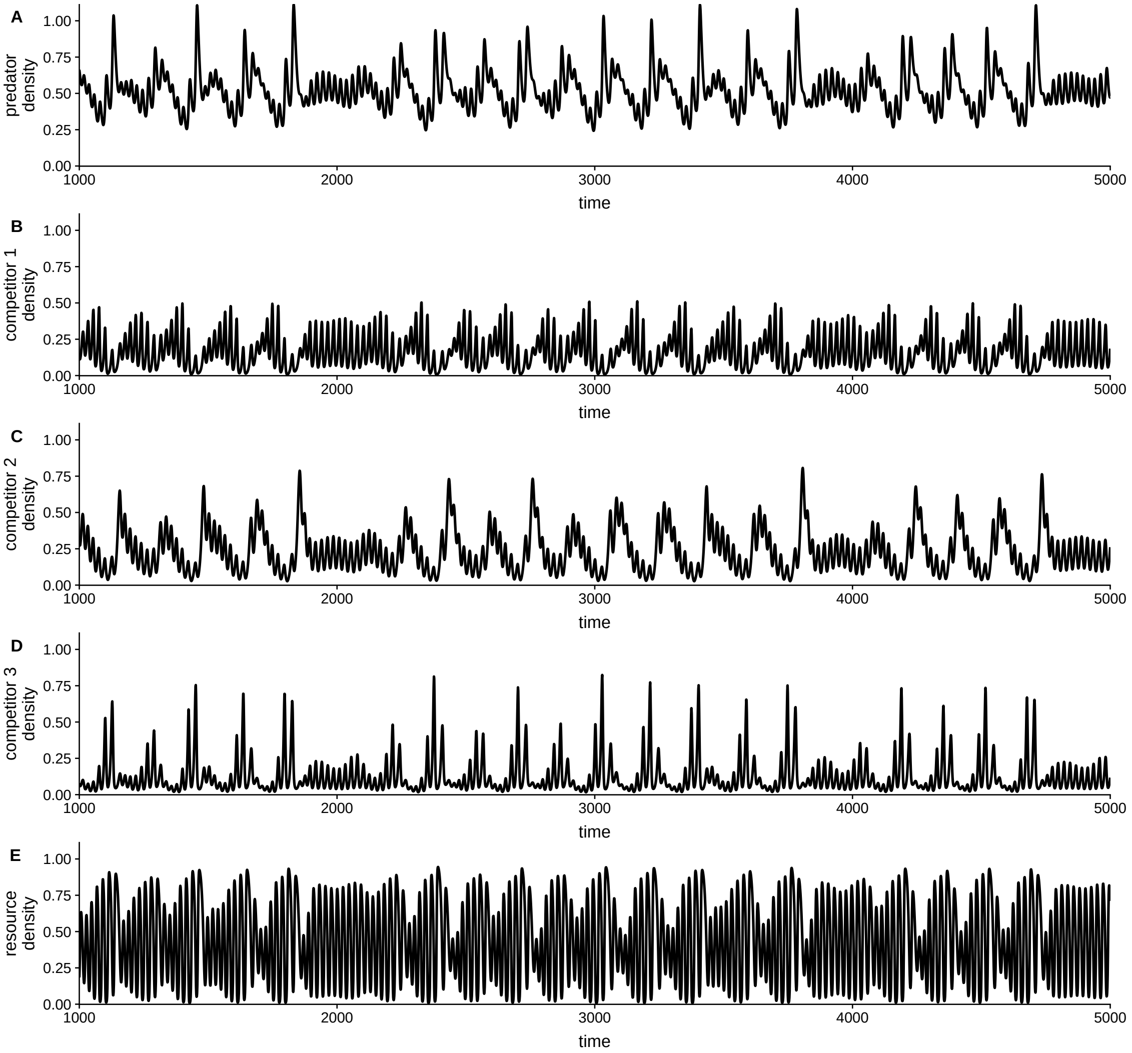


Figure S3.1: Dynamics of the 3 competitor system, from the first set of parameters in the main text (Table 1), as denoted in bold. The figure shows dynamics without environmental perturbations altering mortality rates.

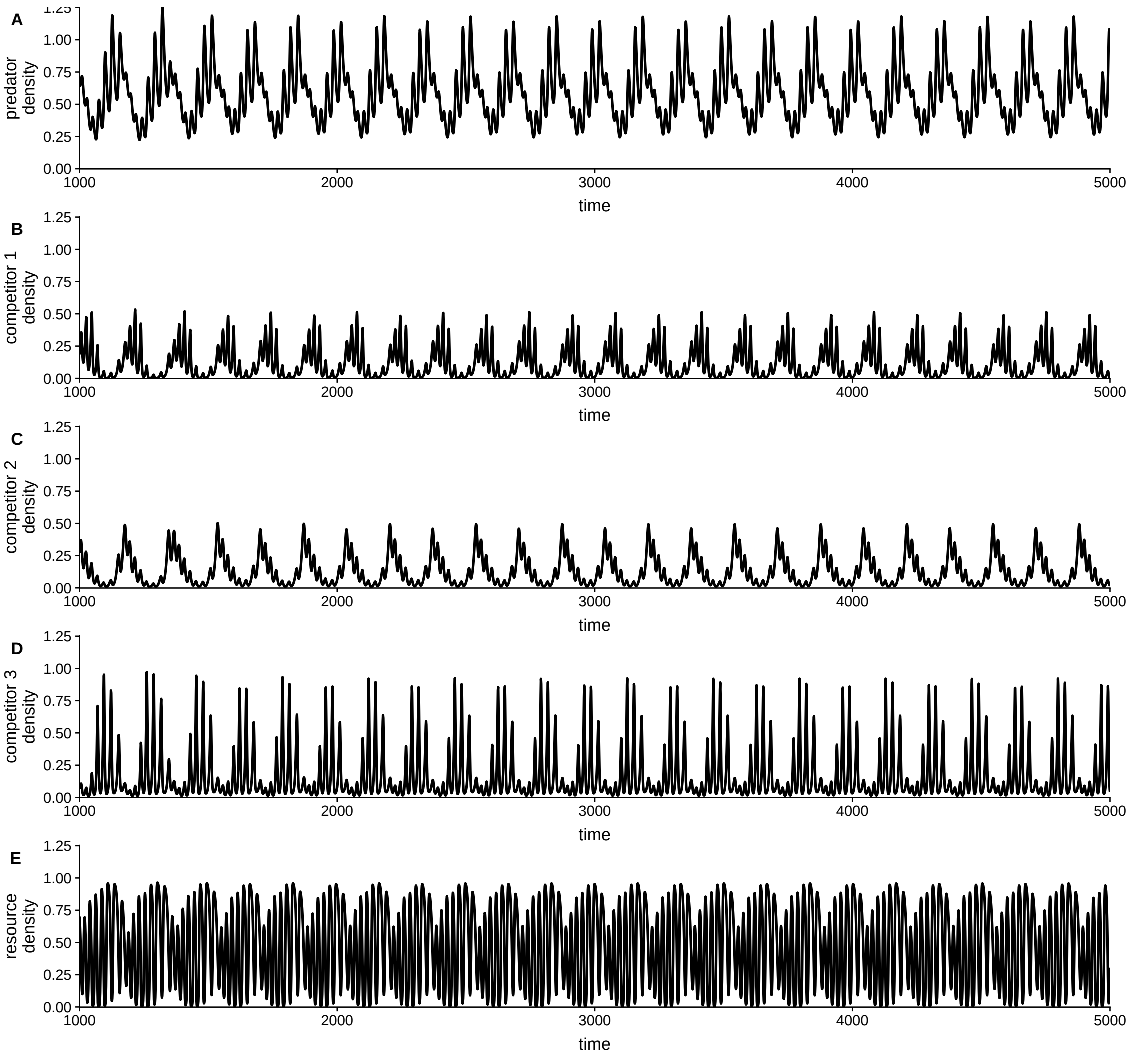


Figure S3.2: Dynamics of a 3 competitor system. These values of the parameters are the second set of parameters given in Table 1. The figure shows dynamics without environmental perturbations altering mortality rates.

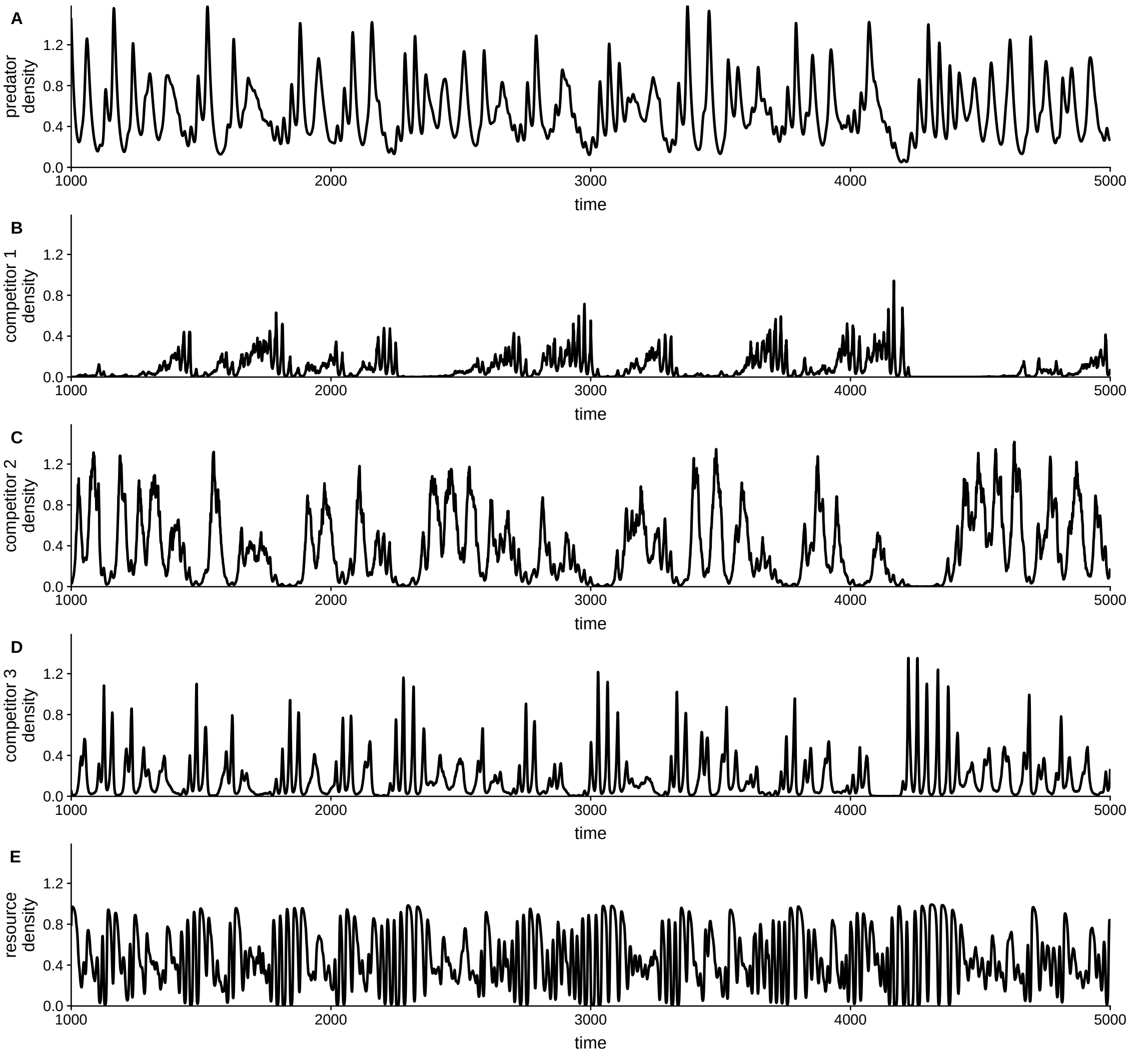


Figure S3.3: Example dynamics of a 3 competitor system (Fig. S3.1) when competitor mortality is impacted by environmental fluctuations.

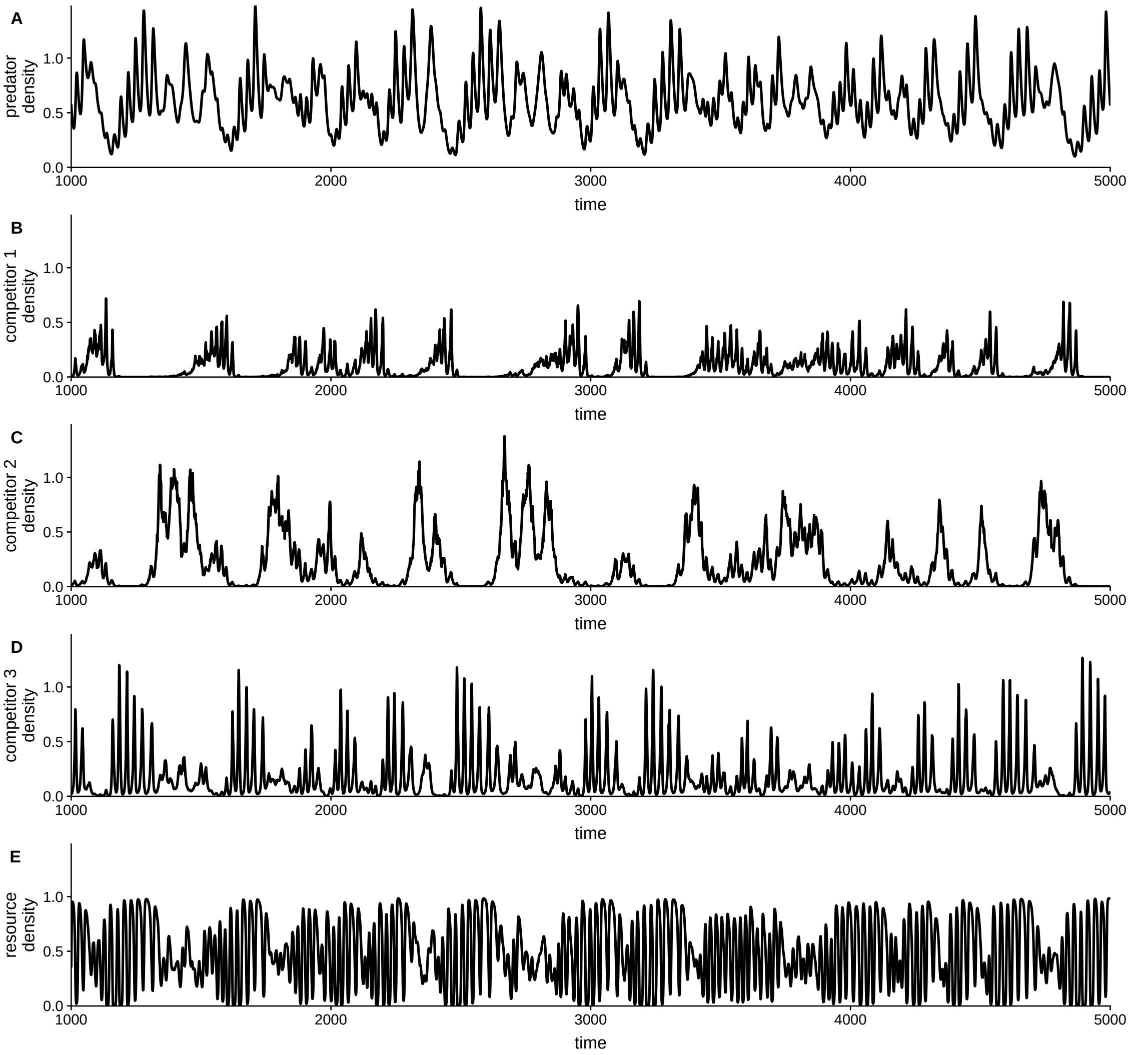


Figure S3.4: Example dynamics of a 3 competitor system (Fig. S3.2) when competitor mortality is impacted by environmental fluctuations.

#### 2 Three Consumers, Two Predators

Expanding the model further to include a second predator yields:

$$\frac{dP_2}{dt} = -M_{p_2}P_2 + \frac{J_{p_2}P_2[\Omega_{P_2C_1}C_1 + \Omega_{P_2C_2}C_2 + (1-(\Omega_{P_2C_1} + \Omega_{P_2C_2}))C_3]}{\tau}$$

$$\frac{dC_1}{dt} = -M_{C_1}C_1 + \frac{\Omega_{C_1R}J_{C_1}C_1R}{R+R_{0_1}} - \frac{\Omega_{P_1C_1}J_{p1}P_1C_1}{\tau} - \frac{\Omega_{P_2C_1}J_{p2}P_2C_1}{\tau}$$

$$\frac{dC_2}{dt} = -M_{C_2}C_2 + \frac{\Omega_{C_2R}J_{C_2}C_2R}{R+R_{0_2}} - \frac{\Omega_{P_1C_2}J_{p1}P_1C_2}{\tau} - \frac{\Omega_{P_2C_2}J_{p2}P_2C_2}{\tau}$$

$$\frac{dC_3}{dt} = -M_{C_3}C_3 + \frac{\Omega_{C_3R}J_{C_3}C_3R}{R+R_{0_3}} -$$
$$\frac{(1-(\Omega_{P_1C_1} + \Omega_{P_1C_2}))J_{p_1}P_1C_3}{\tau} - \frac{(1-(\Omega_{P_2C_1} + \Omega_{P_2C_2}))J_{p_2}P_2C_3}{\tau}$$

$$\frac{dR}{dt} = rR(1-R/K) - \frac{\Omega_{C_1R}J_{C_1}C_1R}{R+R_{0_1}} - \frac{\Omega_{C_2R}J_{C_2}C_2R}{R+R_{0_2}} - \frac{\Omega_{C_3R}J_{C_3}C_3R}{R+R_{0_3}}$$

*where*

$$\tau = \Omega_{P_1C_1}C_1 + \Omega_{P_1C_2}C_2 + (1-(\Omega_{P_1C_1} + \Omega_{P_1C_2}))C_3 +$$

$$\Omega_{P_2C_1}C_1 + \Omega_{P_2C_2}C_2 + (1-(\Omega_{P_2C_1} + \Omega_{P_2C_2}))C_3 + C_0.$$

(S3.2)

We again confirm that estimated parameters result in stable dynamics in the absence of variability in mortality rates before examining coexistence with both environmental and predator fluctuations.

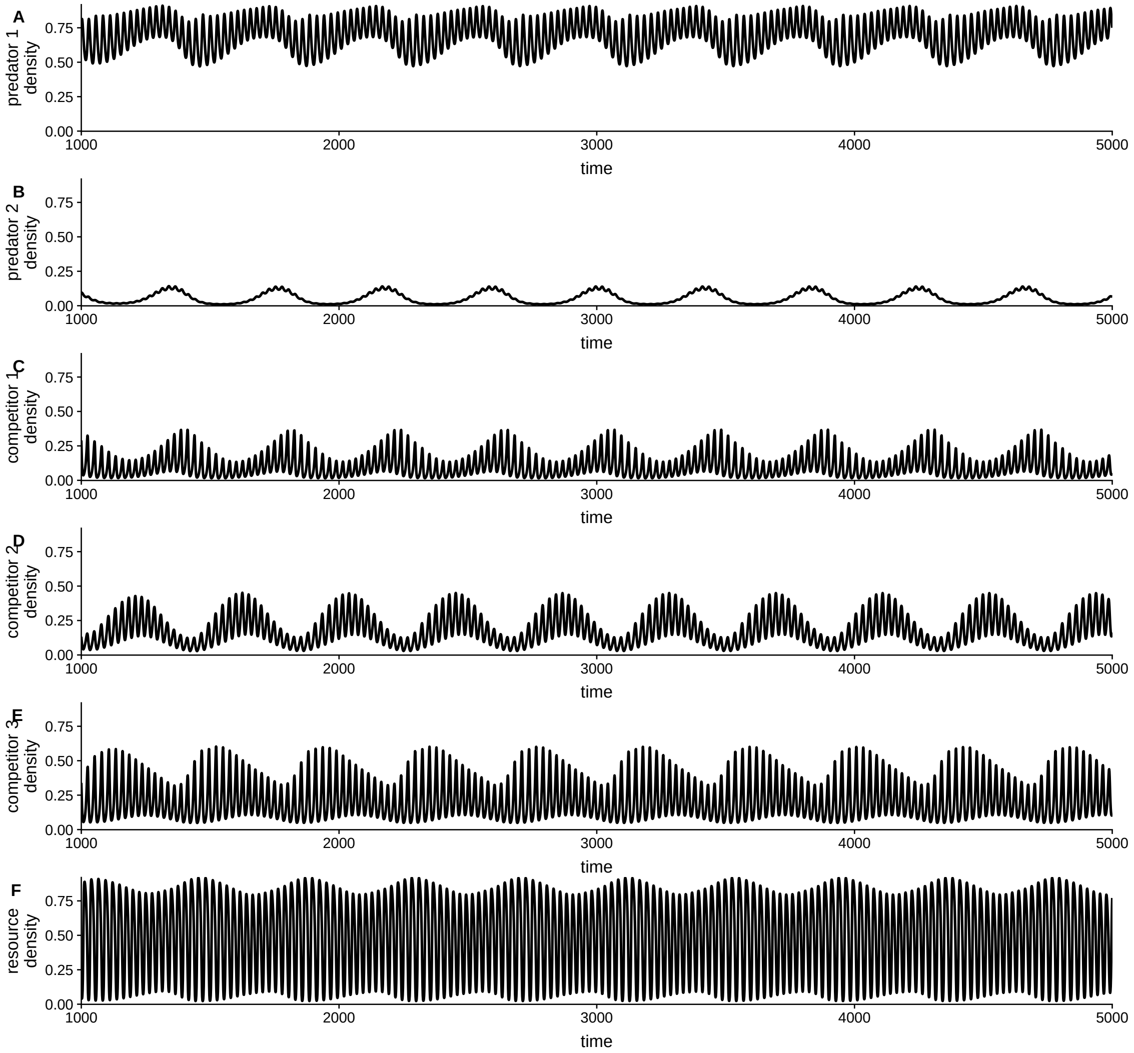


Figure S3.5: Dynamics of the 3 competitor and 2 predator system in the absence of variation in mortality rates. Parameters are given in Table 1 of the main text.

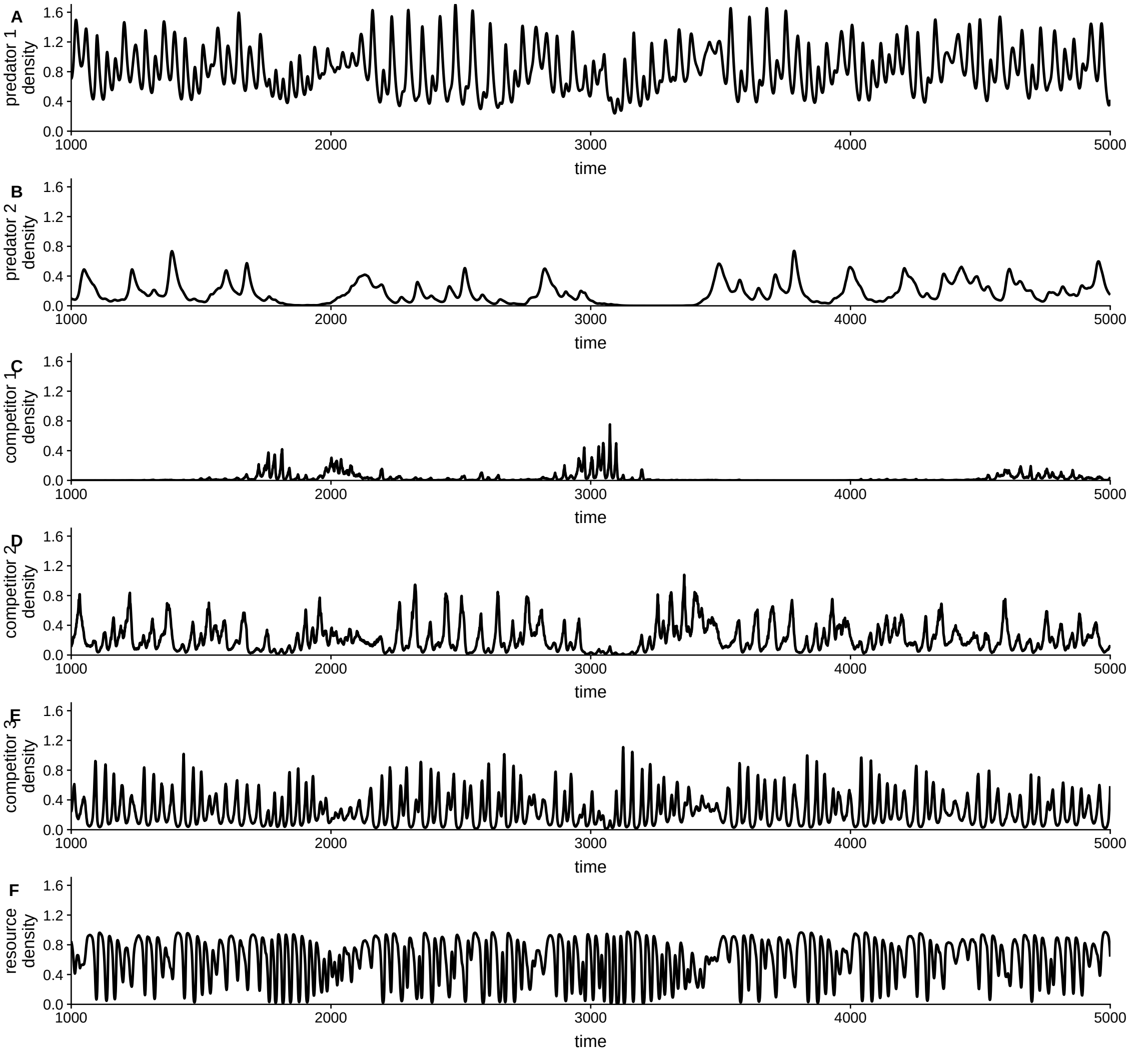


Figure S3.6: Example dynamics of the 3 competitor and 2 predator system with environmental variation. Parameters are given in Table 1 of the main text.

### 3 Coexistence Comparisons

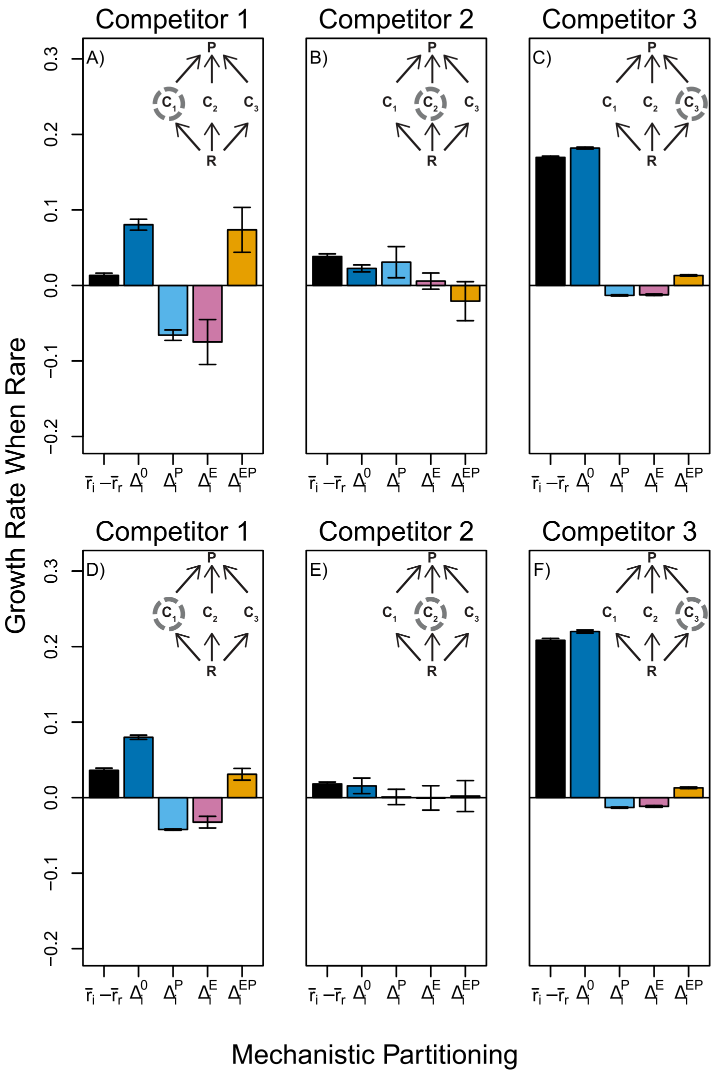


Figure S3.7: Decomposition of the 3 competitor model under two different characterizations. A-C) Three competitor model under the parameter set from replicate 1, as shown in bold in Table 1 of the main text. Data shown here is the same as given in Fig. 5B-D, but is reproduced here for easy comparison with panels D-F. D-F) Three competitor model under the parameter set from replicate 2 given in Table 1 in the main text.
