## Supplement 4 for "Quantifying the relative importance of competition, predation, and environmental variation for species coexistence"

### Rocky Intertidal Food Web Case Study: Supplement 4

Lauren G. Shoemaker, Allison K. Barner, Leonora S. Bittleston, and Ashley I. Teufel

### 1 Model

Our model for rocky intertidal food web dynamics closely builds on the model proposed by Forde & Doak (2004). This model is an extension of foundational work by Iwasa & Roughgarden (1986) and Connolly & Roughgarden (1999). We generally followed the model proposed by Forde & Doak (2004), but made several small changes to facilitate model implementation. Equations S4.3, S4.8, S4.7, and S4.10 were modified from Forde & Doak (2004), described below.

Free space at each time step is a function of the total available space ($T_t$) and the area taken up by the three space-limited competitors (***B. glandula***, ***C. dalli***, and limpets). The total area for each of the three competitor species is proportional to their population size (e.g., $B_t$) and the average size of an adult individual (e.g., $A_b$, Table S4.1). Free space is limited to between 0 and 1 ($m^2$, see Section 2 for more on parameters and units). Free space at time $t$ follows:

$$F_t = T_t - (B_t A_b + C_t A_c + L_t A_l) \tag{S4.1}$$

Barnacles and limpets have pelagic larvae, so recruitment at a given time step (i.e. month) is not related to local abundance and is instead a mass-action process (Connolly and Roughgarden, 1999; Wieters et al., 2008). For each of the these three competitor species, recruitment was thus modeled as a function of available space for the species at the last time point, representing the potential (or maximum) recruitment at a given time step. The number of larvae ($m_x$) in the water column is randomly drawn from a lognormal distribution at each time step (Table 1 in Forde & Doak 2004), but the total recruitment from this larval pool can never be larger than the amount of available space. The size of the larval pool at time $t$ for species $X \subset \{B, C, L\}$:

$$M_{x,t} = \frac{F_{t-1}}{A_x}\left[1 - e^{-a_x m_x / F_{t-1}}\right] \tag{S4.2}$$

Realized barnacle and limpet recruitment was based on Equation 1 in Iwasa and Roughgarden (1986), where settlement of planktonic larvae into the local system depends on the amount of free space at the last time step, the number of larvae of the species in the larval pool (the "potential" recruitment), and the rate of larval settlement of the species (Table S4.1). Actual (realized) recruitment at time $t$ for species $X \subset \{B, C, L\}$:

$$R_{x,t} = d_x M_t F_{t-1} \tag{S4.3}$$

For the two barnacle species, population size at a given time point is a function of adult survivorship from the previous time step, density-independent survival of recruits from the larval pool, and the loss of adults to sea star and whelk predation. Following the assumption of Forde & Doak (2004), recruits transition into the adult population after one month. Population size for barnacle species $Y \subset \{B, C\}$ at time $t$:

$$Y_t = Y_{t-1} S_y + s_y R_{y,t} - p_{whelk} W_{t-1} Y_{t-1} S_y - p_{seastar} P_{t-1} Y_{t-1} S_y \tag{S4.4}$$

The Forde & Doak limpet population model, unlike barnacles, does not include explicit mortality due to predation. Instead, predation on the limpet population is implicitly modeled as density dependence. Thus, limpet population size is a function of adult survival from the previous time step, and density-dependent survival of recruits from the larval pool. Density-dependence, following Forde & Doak (2004), is given by a parameter $\delta$ (Table S4.1). According to Forde & Doak, delta was used to model the density-dependence that would occur if limpet predators were included in the model, but no justification for the parameter value was given. Future work could explicitly include predator dynamics, as in NE Pacific intertidal systems, limpets are eaten by surfperch (Mercurio et al., 1985), birds (Marsh, 1986), sea stars (Phillips and Castori, 1982), and crabs (Lowell, 1986). Similar to barnacles, limpet larval supply is modeled using random draws from a lognormal distribution (Table 1 in Forde & Doak 2004). Population size for limpets at time $t$:

$$L_t = S_l\, L_{t-1} + s_l\, R_{l,t}\, e^{\delta\, L_{t-1}} \tag{S4.5}$$

Whelks lay egg masses once per year, unlike all other species in the model with planktonic larvae. As such, recruitment was calculated once a year (June), and modeled as follows from Forde & Doak (2004). The potential number of new whelk recruits in June was a function of the barnacle prey consumed in the previous three months. $C_{average}$ and $B_{average}$ are the average number of barnacle prey available in April to June, $p_w$ is whelk predation rate, and $\gamma$ is the predator conversion rate. Thus, potential whelk recruitment at time $t$:

$$M_{w,t} = (C_{average} + B_{average})\, 3\, p_{whelk}\, \gamma\, S_{w,t}\, (C_{t-1} + B_{t-1}) \tag{S4.6}$$

In the original Forde & Doak model (2004), a step function was used to implement density-dependence in whelk recruitment (e.g., if $R_w > 90$, then set $R_w$ = 90). Here, we instead used a discrete logistic population equation, with a carrying capacity ($K$) of 90. This required an additional assumption of the population growth rate ($r$, Table S4.1). Actual (realized) whelk recruitment at time $t$:

$$R_{w,t} = M_{w,t}\, \frac{M_{w,t}}{K_w}\, e^{r_w(1-\frac{M_{w,t}}{K_w})} \tag{S4.7}$$

To model whelk adult population size, we simplified the equation from Forde & Doak 2004, so that the whelk population size is modeled similarly to that of the other predator in the system (sea stars; Connolly and Roughgarden (1999)). Population size is simply a function of adults that survived from the previous month (with constant per capita mortality) and new whelk recruits (if June). Population size for whelks at time $t$:

$$W_t = W_{t-1} * S_w + R_{w,t} \tag{S4.8}$$

The larval pool for predator sea stars (*Pisaster ochraceus*), like barnacles and limpets, was modeled as random draws from a lognormal distribution (Table 1 in Forde & Doak 2004). However, sea stars are assumed to not compete for space with barnacles and limpets in this model. Thus, sea star recruitment is simply a function of the larval pool in the water column at a given point in time, and is not related to the available free space. Sea star recruitment at time $t$:

$$\ln(R_{p,t}) \sim \mathcal{N}(\mu_p, \sigma_p^2) \tag{S4.9}$$

Similar to the modified whelk recruitment equation, the population model for sea star abundance was modified from Forde & Doak 2004 to explicitly incorporate density dependence. The step function of Forde & Doak (if $P_t < 6$, $P_t$ = $P_t$, else $P_t$ = 6) was updated to saturate at a carrying capacity of 6, with population size a function of adult survival at the last time point and recruitment into the system. Note that unlike whelk population size, sea star populations are not a function of local prey abundance (Wieters et al., 2008) because adult sea star abundance is instead related to the size of the regional larval pool (Connolly and Roughgarden, 1999). Sea star population size at time $t$:

$$P_t = \rho_t\, \frac{\rho_t}{K_p}\, e^{r_p(1-\frac{\rho_t}{K_p})} \tag{S4.10}$$

Where, $\rho_t = (S_p P_{t-1}) + (s_p R_p)$.

### 2 Parameterization

When possible, we used the parameters from Forde & Doak (2004) Tables 1 and 2; all parameters came from: Burrows and Hughes (1991); Forde (2002); Forde and Doak (2004); Frank (1965); Menge et al. (1994), and Palmer (1990) (Table S4.1). We modified the realized recruitment equation for barnacles and limpets (equation S4.3), which included a new per capita settlement parameter, $d$. Following Connolly & Roughgarden (1999), the settlement coefficient was the same for all three species (see also Gilman (2006) for independent derivation of settlement rate for limpets). Further, the original Forde & Doak model did not include survival rates for sea star recruits. We calculated recruit survival from Menge (1975), given two pieces of information: the average annual survival of spawned gametes to postmaturity longevity is 1.46 x $10^{-9}$/$m^2$/year and the annual mortality of gametes is 0.999 Menge (1975). Whelk and sea star population models were rewritten from a step function in Forde & Doak (2004) to a density-dependent logistic form. To do so, we simply assumed the population growth rate of sea stars was 1, while the whelk population growth rate was much lower. The lower whelk population growth rate was set lower (0.3) after preliminary runs of the model found that a growth rate of 1 resulted in very strong density-dependence that held the whelk population size at fewer than 1—well below the initial population size of 93.

We used the same barnacle adult and recruit survival rates as Forde & Doak (2004). Future work could explore model dynamics if *Balanus* survives at a higher rate than *Chthamalus*, as has been empirically shown in Connell (1961).

Similarly, we continue to use the model form of Forde & Doak such that neither predator has a prey preference, although past experiments suggest that *Balanus* is predated at a higher rate than *Chthamalus* (Connell, 1961; Navarrete et al., 2000). Instead, we increased the per-capita rate of predation to mimic the effect of strong predation on barnacles. Finally, the initial population size was set to be the same for both barnacle species, and was set to the lower initial population size given for *Balanus* in Forde & Doak (2004). Thus, the only difference in the model for the two barnacle species is adult size (*Balanus* $>$ *Chthamalus*) and larval supply rates under "high" supply scenarios (also *Balanus* $>$ *Chthamalus*).

Larval supply rates for barnacles, *Pisaster ochraceous*, and limpets were identical to those in Table 1 in Forde & Doak (2004). For each species, there were three mean values (low, medium, high) and for each mean value, there were three variance values (low, medium, high). For all simulations, we used the "low" variance option that was associated with each of the "low" and "high" mean larval supply values.

| Parameter | Description | Value | Source |
|---|---|---|---|
| *Balanus glandula (B)* | | | |
| $B_0$ | Initial population size | 4100 $m^{-2}$ | Forde (2002) |
| $S_B$ | Adult survival rate | 0.7 $mo^{-1}$ | Connolly & Roughgarden (1999) |
| $S_b$ | Recruit survival rate | 0.7 $mo^{-1}$ | Connolly & Roughgarden (1999) |
| $A_B$ | Average adult size | 0.98 $cm^{-2}$ | Forde (2002) |
| $A_b$ | Average recruit size | 0.03 $cm^{-2}$ | Forde (2002) |
| $d_b$ | Larval settlement coefficient* | 1.44 $mo^{-1}$ | Connolly & Roughgarden (1999) |
| *Chthamalus fissus/dalli (C)* | | | |
| $C_0$ | Initial population size | 4100 $m^{-2}$ | Forde (2002) |
| $S_C$ | Adult survival rate | 0.7 $mo^{-1}$ | Connolly & Roughgarden (1999) |
| $S_c$ | Recruit survival rate | 0.7 $mo^{-1}$ | Connolly & Roughgarden (1999) |
| $A_C$ | Average adult size | 0.32 $cm^{-2}$ | Forde (2002) |
| $A_c$ | Average recruit size | 0.03 $cm^{-2}$ | Forde (2002) |
| $d_c$ | Larval settlement coefficient* | 1.44 $mo^{-1}$ | Connolly & Roughgarden (1999) |
| Limpets (L) | | | |
| $L_0$ | Initial population size | 239 $m^{-2}$ | Forde (2002) |
| $S_L$ | Adult survival rate | 0.97 $mo^{-1}$ | Frank (1965) |
| $S_l$ | Recruit survival rate | 0.88 $mo^{-1}$ | Frank (1965) |
| $A_L$ | Average adult size | 0.8 $cm^{-2}$ | Forde (2002) |
| $A_l$ | Average recruit size | 0.03 $cm^{-2}$ | Forde (2002) |
| $d_l$ | Larval settlement coefficient* | 1.44 $mo^{-1}$ | Connolly & Roughgarden (1999) |
| $\delta$ | Density-dependent paramater | -0.02 | Forde & Doak (2004) |
| Whelks (W) | | | |
| $W_0$ | Initial population size | 93 $m^{-2}$ | Forde (2002) |
| $r_w$ | Population growth rate of whelk recruits* | 0.3 | |
| $S_W$ | Adult survival rate | 0.94 $mo^{-1}$ | Burrows & Hughes (1991) |
| $S_w$ | Recruit survival rate* | 0.88 $mo^{-1}$ | |
| $K_W$ | Carrying capacity* | 90 | Forde & Doak (2004) |
| $p_{W,B}$ | Per capita predation rate on *Balanus** | 0.02 | Forde & Doak (2004) |
| $p_{W,C}$ | Per capita predation rate on *Chthamalus** | 0.02 | Forde & Doak (2004) |
| $Y_W$ | Conversion rate | 0.001 | Forde & Doak (2004) |
| *Pisaster ochraceus (P)* | | | |
| $P_0$ | Initial population size | 1 $m^{-2}$ | Menge et al. (1994) |
| $r_p$ | Population growth rate of adult sea stars* | 1.0 | |
| $S_P$ | Adult survival rate | 0.992 $mo^{-1}$ | Connolly & Roughgarden (1999) |
| $S_w$ | Recruit survival rate* | 0.001 $mo^{-1}$ | Menge (1975) |
| $K_W$ | Carrying capacity* | 6 | Forde & Doak (2004) |
| $p_{W,B}$ | Per capita predation rate on *Balanus** | 0.02 | Forde & Doak (2004) |
| $p_{W,C}$ | Per capita predation rate on *Chthamalus** | 0.02 | Forde & Doak (2004) |

Table S4.1: Parameters used in the model. Descriptions marked with an asterisk indicate parameters that were not included in or modified from the original Forde & Doak (2004) model.

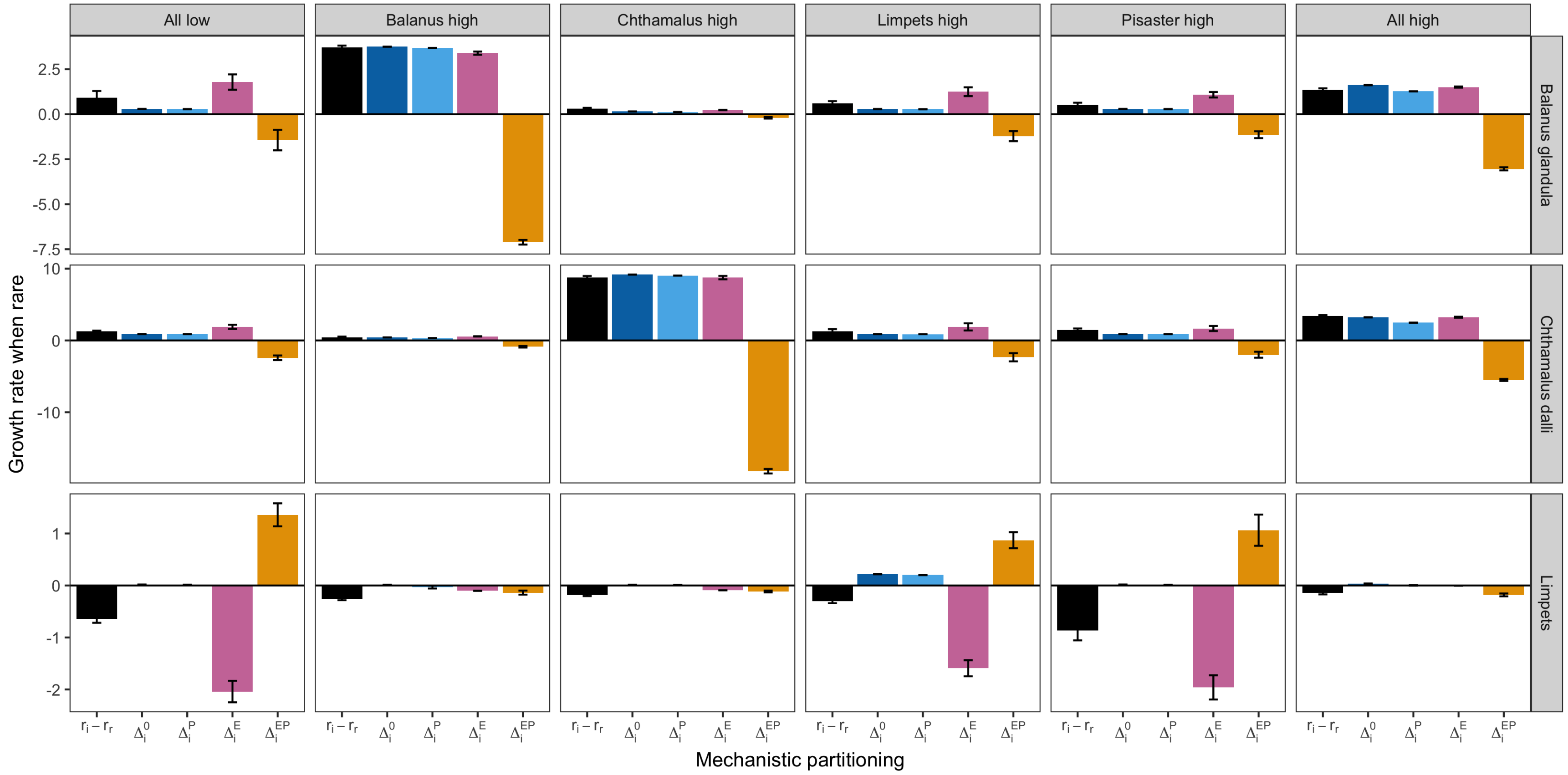


Figure S4.1: Application of coexistence partitioning to the empirical intertidal model, across all levels in larval supply. From the main text: all species have high or low supply, then individually each species has high larval supply while others have low supply. Results show mean and standard error across 500 replicates, each run for 100 years.

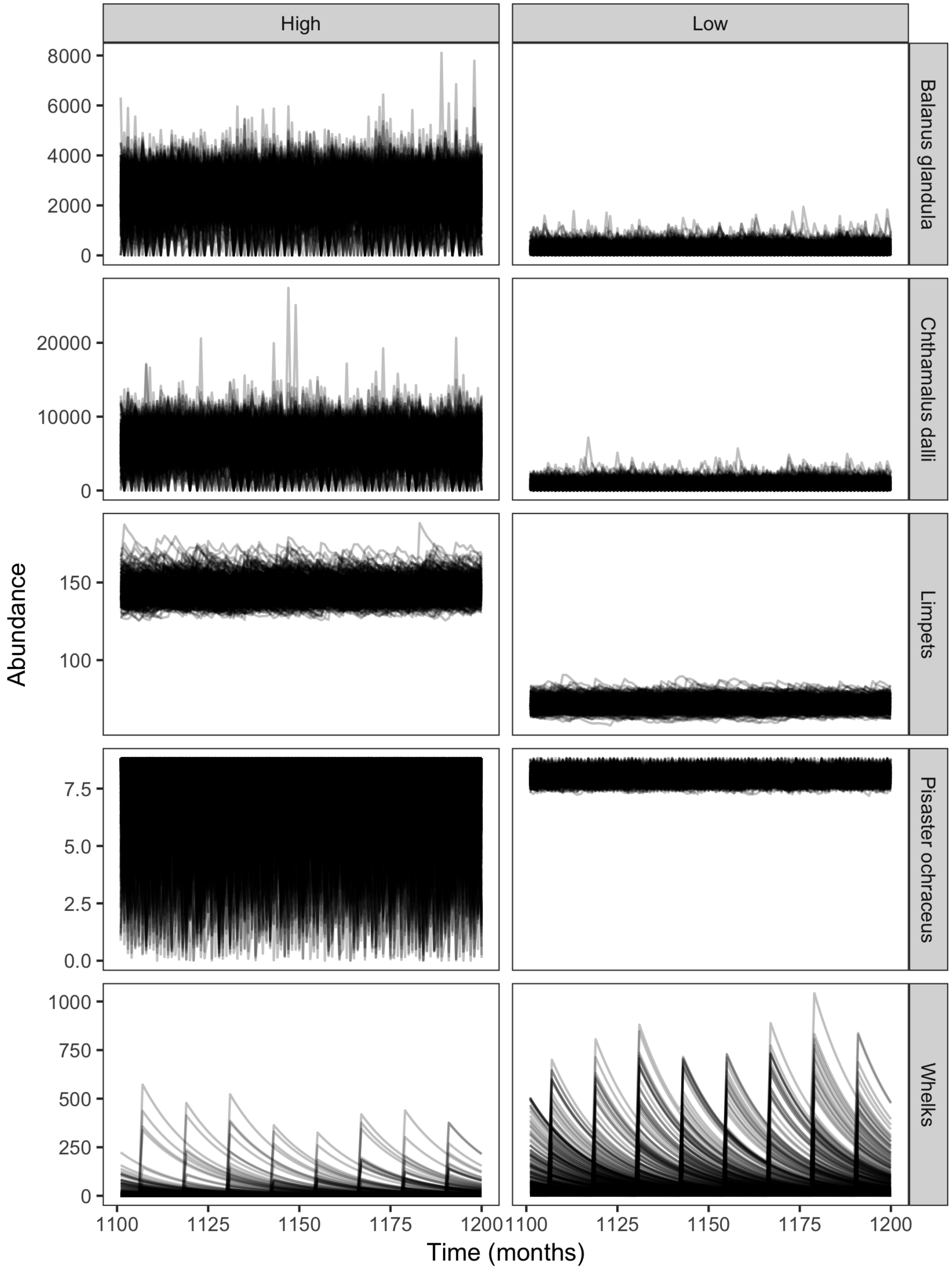


Figure S4.2: Last 100 time steps (months) for 500 runs of the intertidal model, under ”low” and ”high” larval supply. Here, all species were started at their initial population size given in Table S4.1 and then run with all species having either ”low” or ”high” larval supply, as given in Forde & Doak Table 1.
